# High-throughput antibody engineering in mammalian cells by CRISPR/Cas9-mediated homology-directed mutagenesis

**DOI:** 10.1101/285015

**Authors:** Derek M Mason, Cédric R Weber, Cristina Parola, Simon M Meng, Victor Greiff, William J Kelton, Sai T Reddy

## Abstract

Antibody engineering is performed to improve therapeutic properties by directed evolution, usually by high-throughput screening of phage or yeast display libraries. Engineering antibodies in mammalian cells offers advantages associated with expression in their final therapeutic format (full-length glycosylated IgG), however, the inability to express large and diverse libraries severely limits their potential throughput. To address this limitation, we have developed homology-directed mutagenesis (HDM), a novel method which extends the concept of CRISPR/Cas9-mediated homology-directed repair (HDR). HDM leverages oligonucleotides with degenerate codons to generate site-directed mutagenesis libraries in mammalian cells. By improving HDM efficiency (>35-fold) and combining mammalian display screening with next-generation sequencing (NGS), we validated this approach can be used for key applications in antibody engineering at high-throughput: rational library construction, novel variant discovery, affinity maturation, and deep mutational scanning (DMS). We anticipate that HDM will be a valuable tool for engineering and optimizing antibodies in mammalian cells, and eventually enable directed evolution of other complex proteins and cellular therapeutics.

## INTRODUCTION

Following their initial discovery, antibody drug candidates typically require further engineering to increase target affinity or improve a number of other characteristics associated with therapeutic developability (e.g., immunogenicity, stability, solubility)^1^. This is independent of the original source of the antibody (i.e., immunized animals, recombinant or synthetic libraries)^2^. Even with a lead candidate to start from, the potential protein sequence space to explore and optimize for all the relevant drug parameters expands astronomically. Therefore, antibody engineering is done at high-throughput by library mutagenesis and directed evolution using surface display screening, most notably phage and yeast display^3–6^. With some exceptions^7,8^, these display systems typically express antibody proteins as fragments [e.g., single-chain fragment variable (scFv) and fragment antigen binding (Fab)] and without certain post-translational modifications (i.e., glycosylation). However, for therapeutic production, scFvs and Fabs require conversion into full-length glycosylated IgG molecules which consequentially leads to a final optimization phase of evaluating and modifying drug candidates directly in mammalian cells. This step is performed at low-throughput due to the challenges associated with generating libraries in mammalian systems (i.e., inability to stably retain and replicate plasmids).

When engineering candidate antibodies, libraries are often constructed by PCR mutagenesis (e.g., error-prone PCR and site-directed mutagenesis with degenerate primers), followed by cloning into expression plasmids, making them compatible for screening by phage and yeast display. With the motivation of being able to screen antibodies in their native context as full-length IgGs with proper glycosylation, attempts have also been made to incorporate libraries into mammalian cells using episomal-, viral-, or transposon-mediated gene transfer^9–11^. However, relative to phage (>10^10^) and yeast (>10^7^), these mammalian display systems are substantially challenged by small library size (∼10^4^ variants for genome-integrated libraries) and polyclonality (multiple antibody variants per cell). Therefore, in order to truly have a competitive platform for mammalian antibody engineering, an alternative method which overcomes these limitations is essential.

With the rapid advancements in genome editing technologies, most notably the CRISPR/Cas9 system (Cas9), it is now possible to easily make targeted genomic modifications in mammalian cells^12^. While Cas9 is most widely used for gene knock-out (via non-homologous end joining, NHEJ) or gene knock-in (via HDR), it also enables the generation of libraries in mammalian cells. For example, Cas9 has been used to promote HDR with degenerate templates, resulting in a library of genomic variants; this has been applied to both coding and non-coding regions, providing insight into gene regulation, expression, and even drug resistance^13,14^. In a recent study, Cas9 was also used to integrate a genomic landing pad containing a recombination site, which allowed for the introduction of a library of transgene variants^15^. Although these studies illustrate the potential to integrate libraries into specific genomic regions of mammalian cells, transfection of genome editing reagents combined with low HDR efficiencies limit the scalability and ease-of-use required to generate libraries capable of exploring sufficient protein sequence space, which is crucial for directed evolution and protein engineering.

In this study, we have established the method of HDM, which relies on high-efficiency HDR by Cas9 to generate site-directed mutagenesis libraries in mammalian cells. We use as our mammalian antibody display platform, a recently developed hybridoma cell line, where antibody variable regions can be exchanged by Cas9-driven HDR, referred to as plug-and-(dis)play hybridomas (PnP)^16^. A critical feature of our HDM method is that it utilizes single-stranded oligonucleotides (ssODNs) as the donor template, which relative to double-stranded DNA, drastically increase HDR integration efficiencies^17–19^ and also reduce off-target integration events^20,21^. By using a cellular genotype-phenotype assay, we optimized a series of parameters, allowing us to achieve a nearly ∼35-fold improvement of HDR efficiency. Next, starting with an antibody specific for a model antigen, we introduce a library into the variable heavy chain (V_H_) complementarity determining region 3 (CDRH3) by using ssODN templates with commonly used NNK and NNB degenerate codon regions. Following HDM, we perform NGS on the entire V_H_ region to quantitatively assess the library diversity and distribution. We further maximize the efficacy of HDM libraries by implementing an optimized gRNA target sequence and rationally selecting degenerate nucleotides to match target amino acid (a.a.) frequencies of CDRH3 regions found in the antibody repertoire of murine naïve B cells^22,23^. With HDM, we were able to achieve a library size of >10^5^ variants validated by NGS. We then screen this library for specificity towards our model antigen, which led to recovery of a novel variant with a unique CDRH3. We also show that by using HDM to generate saturation libraries across the CDRH3, we could perform directed evolution and affinity maturation.

Finally, we apply HDM to the recently established method of DMS^24,25^, which allowed us to deconstruct the antigen-binding sequence landscape of our antigen-specific antibodies. Through HDM we have successfully developed a rapid and facile method for the generation of site-directed mutagenesis libraries in mammalian cells, which represents a versatile approach for high-throughput antibody engineering.

## RESULTS

### Optimizing HDM efficiency

The degree of genetic diversity and library size that can be introduced using Cas9 is dependent on HDR efficiency. Previously, when establishing our PnP hybridoma platform, we observed an HDR efficiency of less than 1.0% when exchanging a fluorescent reporter protein with antibody variable regions encoded on a double-stranded DNA cassette (∼1.5 kb donor regions, ∼0.7 kb each for left and right homology arms)^16^. Previous studies have shown that despite having only (micro)homology arms, much higher HDR efficiencies are observed when using ssODN donor templates^17,26,27^. Due to the length limitation of commercially synthesized ssODNs, the target region of mutation is typically 50-80 nucleotides (nt), with ∼50 nt for each homology arm, which is highly compatible for targeting antibody CDRs for mutagenesis. However, in contrast to the previous approach, where loss of reporter protein and gain of antibody expression could easily be used to detect HDR, detection of HDR with ssODNs templates is more challenging. Therefore, to quantify and subsequently optimize ssODN-based HDR, we first developed a cellular phenotype assay based on antibody expression and antigen binding.

Starting with the PnP-HEL23 hybridoma cell line, which expresses a murine antibody sequence with specificity towards the model antigen hen egg lysozyme antigen (HEL)^16^, we used Cas9 and guide RNA (gRNA) targeting CDRH3 to introduce a frameshift mutation by NHEJ, resulting in the knockout of antibody expression (PnP-HEL23.FI cell line) (**Fig. 1a**). We then designed an ssODN template that encoded the original CDRH3 a.a. but contained silent mutations, thus Cas9-driven HDR could be detected if both antibody expression and specificity for target antigen HEL were restored. If only antibody expression was detected without binding to HEL, we presumed that an insertion/deletion (indel) via NHEJ or micro-homology mediated end joining (MMEJ) had occurred that knocked the antibody sequence back in-frame. Editing efficiencies were measured by flow cytometry following labelling of cells with fluorescently-tagged HEL and anti-IgH (**Fig. 1b**).

**Fig. 1:**
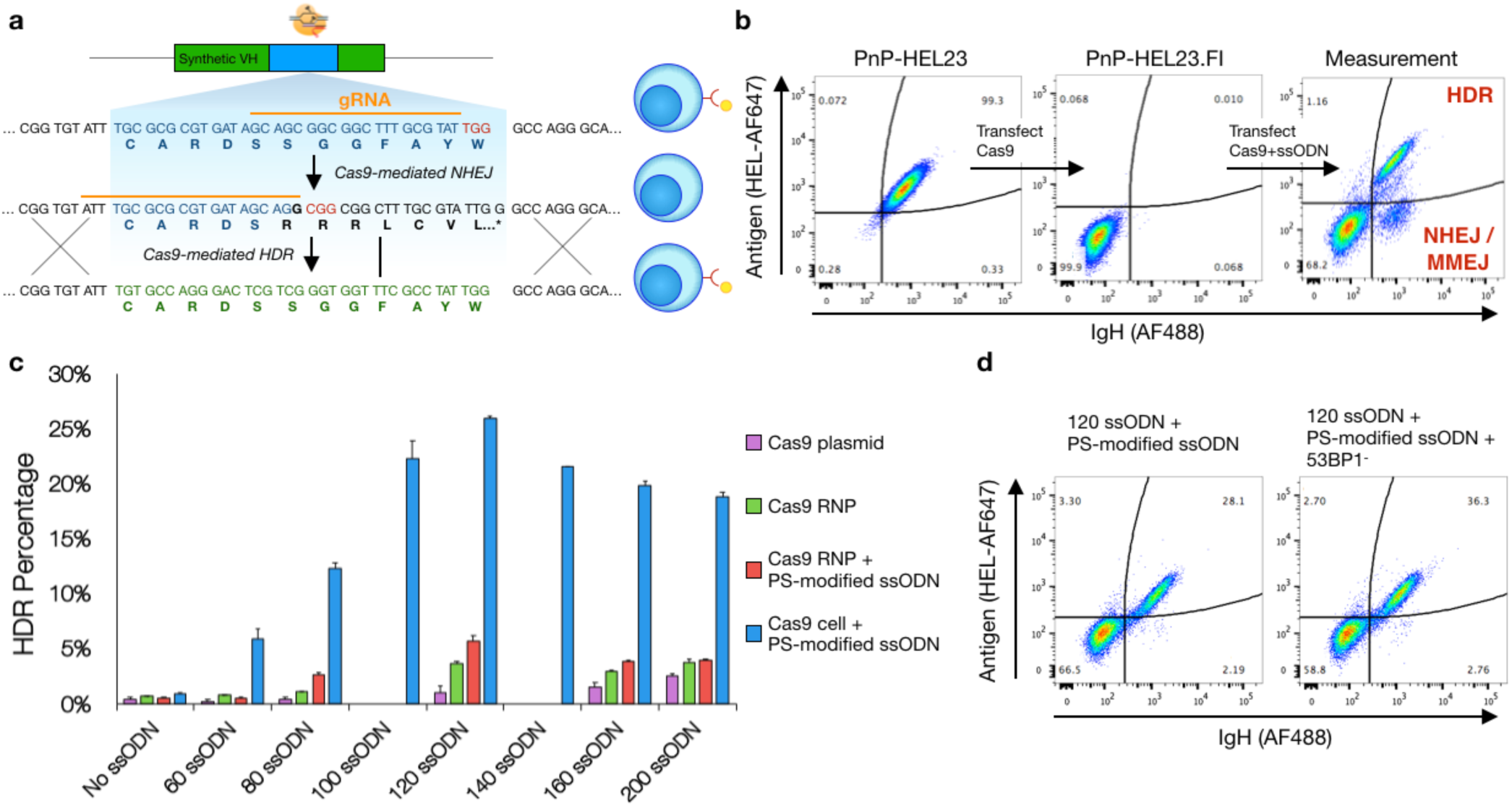
Optimizing parameters for homology directed repair (HDR) **a**, An experimental assay developed to measure HDR efficiencies by flow cytometry. Cas9 is used to knockout antibody expression in the CDRH3 of a PnP hybridoma cell line which originally expressed a functional antibody targeting the model antigen Hen Egg Lysozyme (HEL). A monoclonal population is then isolated and used for all other downstream experiments. Cas9 promotes HDR at the CDRH3 in the presence of a ssODN donor template which contains silent mutations. Cells which undergo HDR will therefore regain antibody expression specific for HEL. **b**, Flow cytometry plots of the original cell line, PnP-HEL23 (left), the CDRH3 knockout cell line, PnP-HEL23.FI (middle), and a representative flow cytometry plot demonstrating the ability of the experimental assay to measure HDR integration efficiencies. **c**, Optimal conditions for HDR revealed by results from assays examining the parameters: i) Cas9 delivery method, ii) ssODN length, and iii) phosphorothioate (PS) modifications to ssODNs, Data presented (mean±sd) is representative of n = 2 independent experiments. **d**, The maximum achievable HDR efficiency of ∼36% was obtained after combing the optimal parameters (constitutive Cas9, ssODN length 120 with PS modifications) and knocking out expression of the 53BP1 protein (PnP-HEL23.FS cells used here, see **Fig. 2**).

To maximize HDR efficiency in PnP cells, several parameters regarding Cas9, gRNA, and donor templates were evaluated by performing separate and parallel transfections (2×10^5^ cells). To resolve the effect of homology arm length on HDR, ssODNs ranged between 60 and 200 nt. To determine if an increase in resistance to nuclease degradation of ssODNs improves HDR, we included phosphorothioate (PS) bonds in the 5’ and 3’ ends. Cas9 and gRNA was delivered to cells by transfection (electroporation) with either plasmid or ribonucleoprotein (RNP) complexes^18,28^. We also generated a PnP stable cell line which constitutively expresses Cas9 from the Rosa26 safe harbor locus (**Supplementary Fig. 1)**, as it permits the transfection of just pre-formed guide RNA (gRNA) and ssODN donor (constant expression of Cas9 has been shown to have no toxic side effects on cells or *in vivo*^29^). Across all conditions, we observed the most notable increase in HDR efficiency when using the constitutive Cas9-expressing cell line, thus all subsequent experiments were performed with constitutive Cas9 expression **(Fig. 1c, Supplementary Fig. 2).** The highest HDR efficiency of 26% was observed when ssODNs with PS modifications and 120 nt length (homology arms equal to 46 nt). This HDR rate remained stable when scaling up the number of cells transfected and only decreased slightly to ∼20% following transfection of 1 × 10^7^ cells (**Supplementary Fig. 3**). Recently, it has been shown that HDR efficiency can be improved by inhibiting the DNA repair regulator protein 53BP1, which promotes NHEJ over HDR^30,31^. Therefore, we used a gRNA to target and knockout 53BP1 in our Cas9-expressing cell line; whereby subsequent testing of HDR in these cells showed an improved HDR efficiency of ∼36% (120 nt and PS-modified ssODNs), a >35-fold improvement relative to 1% (120 nt, Cas9 plasmid, unmodified ssODNs) (**Fig. 1d**).

**Fig. 2:**
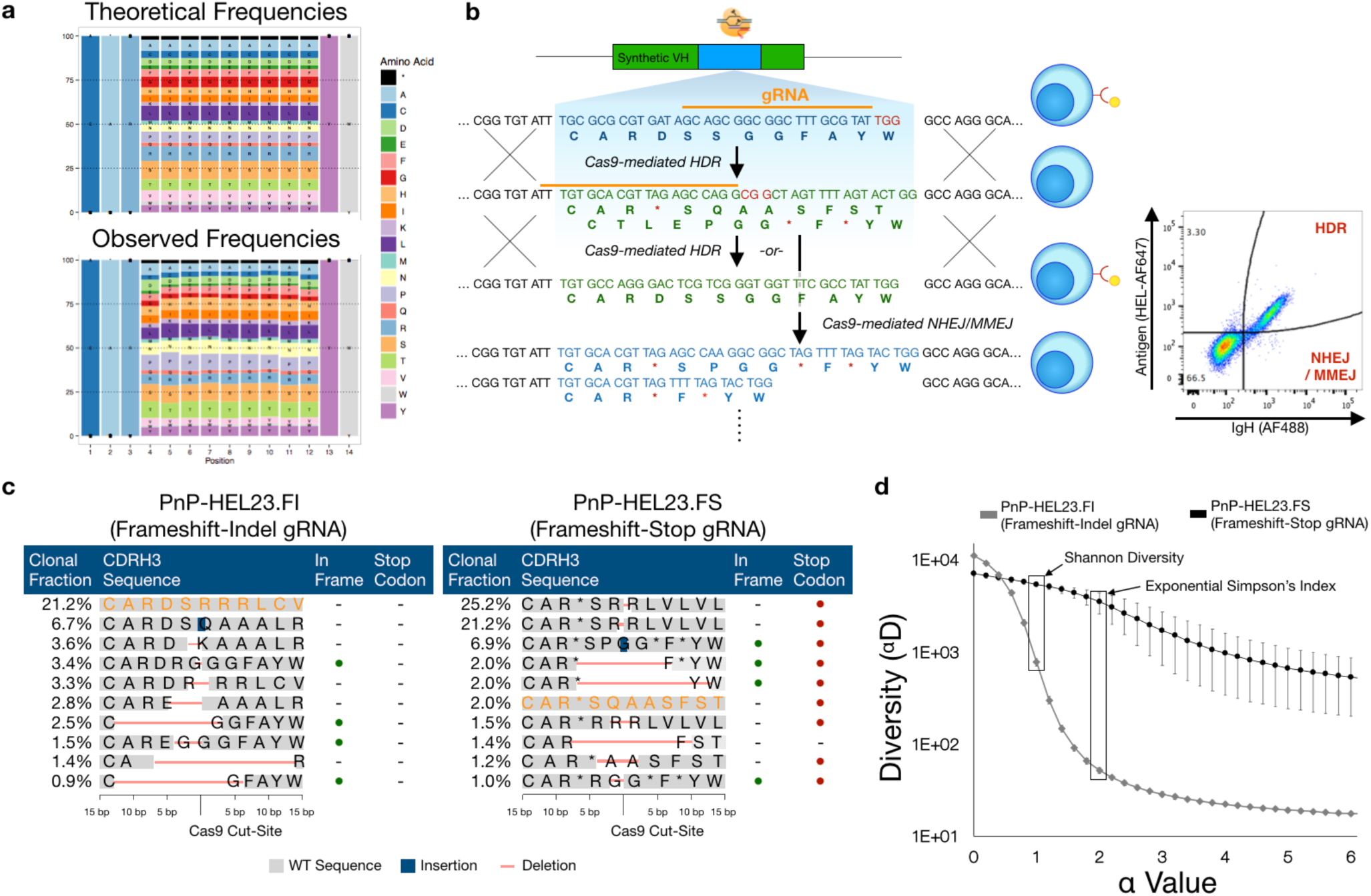
Generating unbiased libraries with homology-directed mutagenesis (HDM) **a**, Amino acid frequencies of observed HDM events when generating NNK/B libraries resembles the theoretical amino acid frequencies, indicating unbiased integration. **b**, A PnP hybridoma cell line was engineered with an optimized, gRNA target sequence to reduce bias arising from NHEJ/MMEJ events. The gRNA sequence adheres to nucleotide propensities observed to increase Cas9 activity6763 while minimizing potential off-target sequences. Following Cas9 cleavage, stop codons encoded on both the 5’ and 3’ ends of the Cas9 cut-site promote recombination of in-frame stop codons following repair via NHEJ/MMEJ. These in-frame stop codons show reduced library bias arising from nonrandom repair. **c**, NGS analysis of the top 10 CDRH3 clones for NNK/B libraries generated from either the frameshift-indel cell line (PnP-HEL23.FI) or the frameshift-stop cell line (PnP-HEL23.FS). In addition to all in-frame NHEJ/MMEJ events containing a stop codon, a much lower percentage of cells matching the originating CDRH3 nucleotide sequence (•) was observed. **d**, Diversity profiles calculated from NGS data for NNK/B libraries generated from the PnP-HEL23.FI or the PnP-HEL23.FS cell lines. The values represent different weights to the clonal frequency, where α = 0 is the species richness, and α = 1 and α = 2 are two common measures for library diversity known as the Shannon entropy and Simpson’s index, respectively. Both the Shannon entropy (α = 1) and Simpson’s Index (α = 2) are higher in the case for libraries generated from the PnP-HEL23.FS cell line, indicating a higher degree of unbiased diversity. Data presented (mean±sd) is representative of n = 4 experiments.

**Fig. 3:**
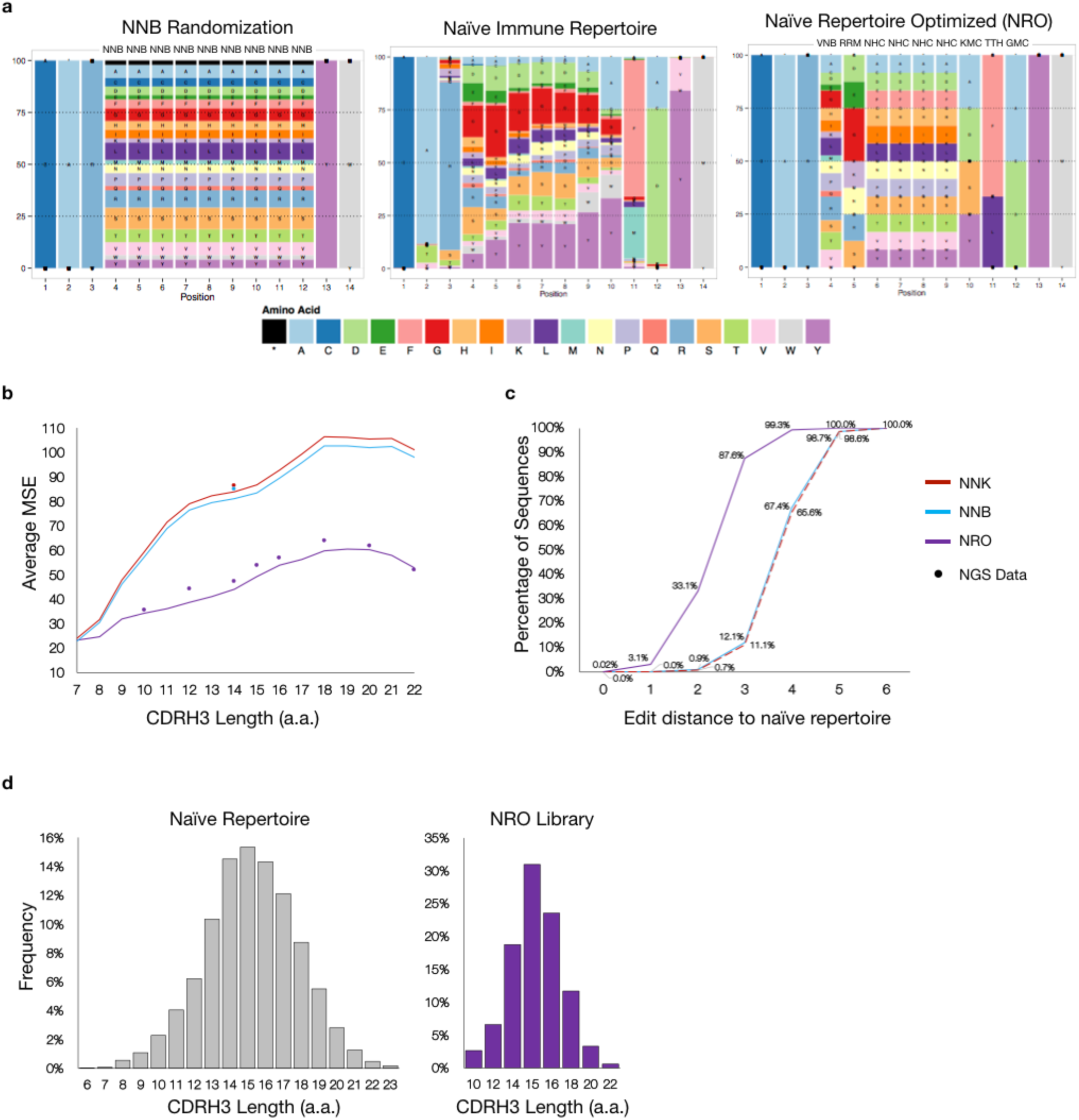
Naïve Repertoire Optimized (NRO) library design and analysis. **a**, Side-by-side comparisons of the CDRH3 amino acid frequencies found in the naïve antibody repertoire (NGS data of naïve B cells from mice) and the theoretical NNB and NRO randomization schemes. **b**, The theoretical and observed averages (from NGS) of the positional mean squared errors (MSEs) evaluated between the naïve repertoire and degenerate codon libraries across various CDRH3 lengths. The NRO scheme more closely resembles the amino acid frequencies observed in the naïve antibody repertoire compared to both NNK and NNB schemes. Equivalent average MSEs for both theoretical and observed amino acid frequencies reflect the accuracy of ssODN synthesis and unbiased HDM integration preferences. **c**, Levenshtein distances for CDRH3 sequences of length 14 between the HDM libraries generated with degenerate codon schemes and the naïve antibody repertoire. The NRO scheme has a higher percentage of CDRH3 sequences with shorter edit distances indicating a higher degree of overall sequence similarity with the naïve repertoire. **d**, Results from NGS data displays the ability to control the CDRH3 length distribution of the HDM library from a single transfection. By accounting for differences in integration efficiencies (**Supplementary Fig. 5**), ssODNs of various lengths were pooled at varying ratios to mimic the length distribution of the naïve antibody repertoire.

### Assessing HDM library diversity and eliminating bias by gRNA design

Targeting the CDRs for site-directed mutagenesis has proven to be an effective means of building antibody libraries for improved properties^32,33^. Introducing sequence diversity into CDRH3 alone has shown to be sufficient for a wide range of antigen-binding specificities^34^. Thus, initial libraries were generated using ssODN homology templates (126 nt) that contained nine consecutive degenerate codons of NNK or NNB within the 14 a.a. long CDRH3 (e.g, ‘*CAR(NNK)*_*9*_*YW’*). These degenerate ssODN donors were transfected with gRNA into PnP-HEL23.FI cells (2 × 10^5^ cells, two replicates per codon scheme). Here, we define the difference between HDR and HDM, in that the latter uses ssODN templates with degenerate codons. In order to precisely quantify library diversity introduced by HDM, following transfection and cell recovery and expansion, genomic DNA was isolated and targeted PCR was performed on the V_H_ region to amplify libraries for NGS (using a previously established protocol for Illumina paired-end sequencing)^35^. Across the four samples of NNK and NNB libraries, sequencing depth ranged between 577,915 – 847,687 read counts, with a ∼95% alignment success, resulting in 12,773 −14,842 unique CDRH3 a.a. sequences (**Supplementary Table 1**). CDRH3 sequences with a length of 14 a.a. were classified as HDM events. Analysis of the HDM sequences revealed nearly an unbiased a.a. usage: positional a.a. usage in HDM libraries was nearly identical to what is theoretically predicted using NNK/B codon scheme (mean-squared error (MSE) = 3.07±0.32) (**Fig. 2a**). Furthermore, overlap analysis of sequences revealed almost no common CDRH3 sequences across all replicates, indicating that each HDM experiment results in a unique library (**Supplementary Table 2**).

In addition to HDM, a subpopulation of cells can restore antibody expression through NHEJ or MMEJ. Consistent with previous reports^36^, we observed that the non-randomness of these pathways created a bias in the library: there were several grossly overrepresented variants (**Fig. 2c**). This presence of high-frequency, redundant variants may limit library diversity and impede screening and selection steps. In order to address this issue, we used the information from the highest frequency NHEJ or MMEJ events to rationally design a gRNA target sequence which would promote either a frameshift mutation or in-frame stop codon following Cas9 cleavage and DNA repair. Using this approach, we drastically reduced the probability that an NHEJ or MMEJ event would result in a functionally expressed antibody by ensuring in-frame stop codons would appear on the 5’ and 3’ sides of the cleavage site (**Fig. 2b**). Next, we used this new, frameshift-stop cell line (PnP-HEL23.FS) to generate HDM libraries, using once again NNK/B ssODN templates. NGS on V_H_ genes was performed as before and similar sequencing depth and quality was obtained (**Supplementary Table 1**). Close analysis of the CDRH3 distribution revealed a substantial decrease in the frequencies of overrepresented (biased) variants and the generation of a more desirable uniformly diverse library. Diversity was quantified using the Hill diversity (**Fig. 2d**, *Equation 2*)^37^, where the diversity (^α^D) for each alpha value represents an equivalent library in which all variants are equally present; Shannon diversity (alpha =1) and Simpson’s index (alpha = 2) are widely used for diversity comparisons.

### Design, generation, and analysis of an HDM library that mimics the naïve antibody repertoire

While we have shown that with optimized parameters we can achieve considerable HDM efficiency, the growth rate and number of mammalian cells that can be handled in a typical experiment still pose challenges in achieving large library sizes (>10^5^). With this in mind, we looked to maximize the functional quality of HDM libraries through the rational selection of degenerate codons. A major limitation in standard NNK/B codon schemes is the compounding probability of introducing a premature stop codon as the number of degenerate codons increases (**Supplementary Fig. 4a**). However, when considering every possible combination of nucleotides, there exist 3,375 different degenerate codon schemes. Thus, as a first approach towards optimizing ssODNs, we designed degenerate codons that would most closely mimic the CDRH3 a.a. frequencies found in the antibody repertoire of murine naïve B cells (433,618 unique CDHR3 sequences)^23^ (**Fig. 3a**). It is well established that CDRH3 libraries with a more natural sequence landscape offer favorable properties (i.e., reduced immunogenicity, improved protein stability and folding)^38^. Therefore, to generate the naïve repertoire optimized (NRO) library, for each position the degenerate codon was selected that produced the minimal MSE value relative to the naïve repertoire, while also using weights to punish degenerate codons that result in cysteines or stop codons (*Equation 1*). Following HDM and NGS, we observed that relative to the naïve repertoire, NRO libraries had substantially lower MSE values (a.a. length 14, average MSE = 47.1) compared to that of NNK/B libraries (a.a. length 14, average MSE = 85.7±0.85) (**Fig. 3b**). Notably, the possibility of introducing a premature stop codon or cysteine residue was eliminated while still maintaining adequate levels of total diversity (**Supplementary Fig. 4b**). Using another indication of sequence similarity, we determined based on the Levenshtein (edit) distance that the NRO library had much higher percentage of sequences with short edit distances compared to NNK/B libraries (**Fig. 3c**).

**Fig. 4:**
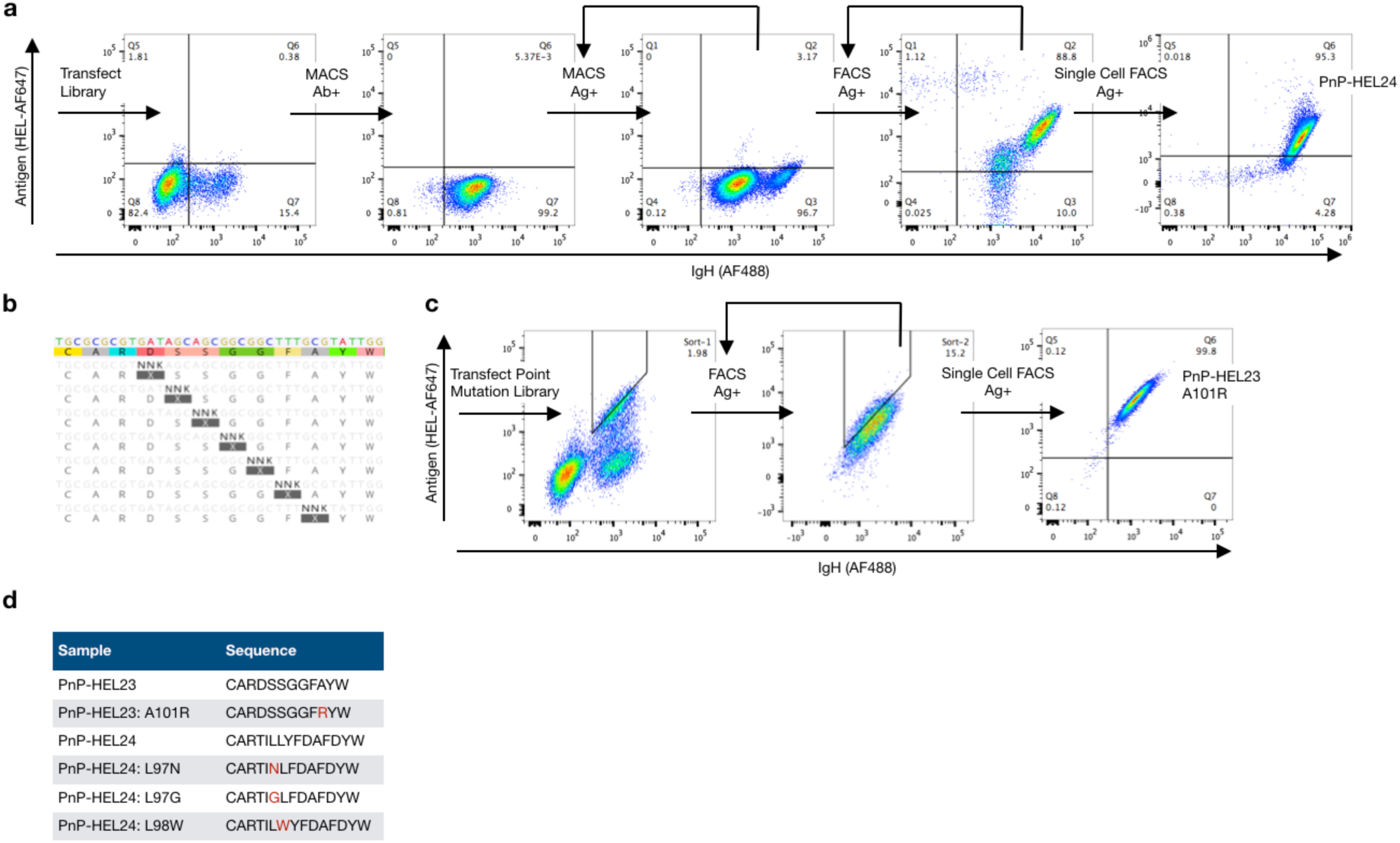
HDM library screening for antibody discovery and affinity maturation. **a**, Library generation by HDM coupled with mammalian cell display screening leads to the discovery of a novel CDRH3 variant, HEL24, for the target antigen, hen egg lysozyme (HEL). Following HDM library generation with NRO ssODNs, antibody expressing cells (Ab+) are isolated by magnetic activated cell sorting (MACS) and expanded. The library is then screened by MACS for specificity to antigen (Ag+), followed by two rounds of screening by fluorescence activated cell sorting (FACS). A monoclonal population for downstream characterization was isolated by single-cell sorting, revealing the novel, variant, HEL24. **b**, Saturation HDM libraries for affinity maturation are created by pooling ssODNs that tile the mutagenesis site along the CDRH3. **c**, Screening of saturation HDM libraries of HEL23 by FACS or higher antigen affinity variants, while using IgG staining to normalize for antibody expression. Monoclonal populations for downstream characterization were isolated by single-cell sorting. **d**, The original HEL23 and HEL24 sequences and their higher affinity variants. ELISA data confirms secretion of antigen specific antibodies into the hybridoma supernatant and antigen affinities relative to one another (**Supplementary Fig. 6**).

The CDRH3 length distribution of natural antibody repertoires also represents another important aspect to recapitulating functional diversity. Therefore, in order to assess the impact of CDRH3 length on HDM efficiency, separate transfections of 2×10^5^ cells were performed with ssODN donors containing various lengths of degenerate codon regions, while keeping homology regions constant. As hypothesized, a minor decrease in integration efficiency was observed as the a.a. degenerate region increases (**Supplementary Fig. 5**). This relationship between degenerate codon region length and integration efficiency can be taken into consideration when building libraries that aim to resemble the natural CDRH3 length distribution.

**Fig. 5:**
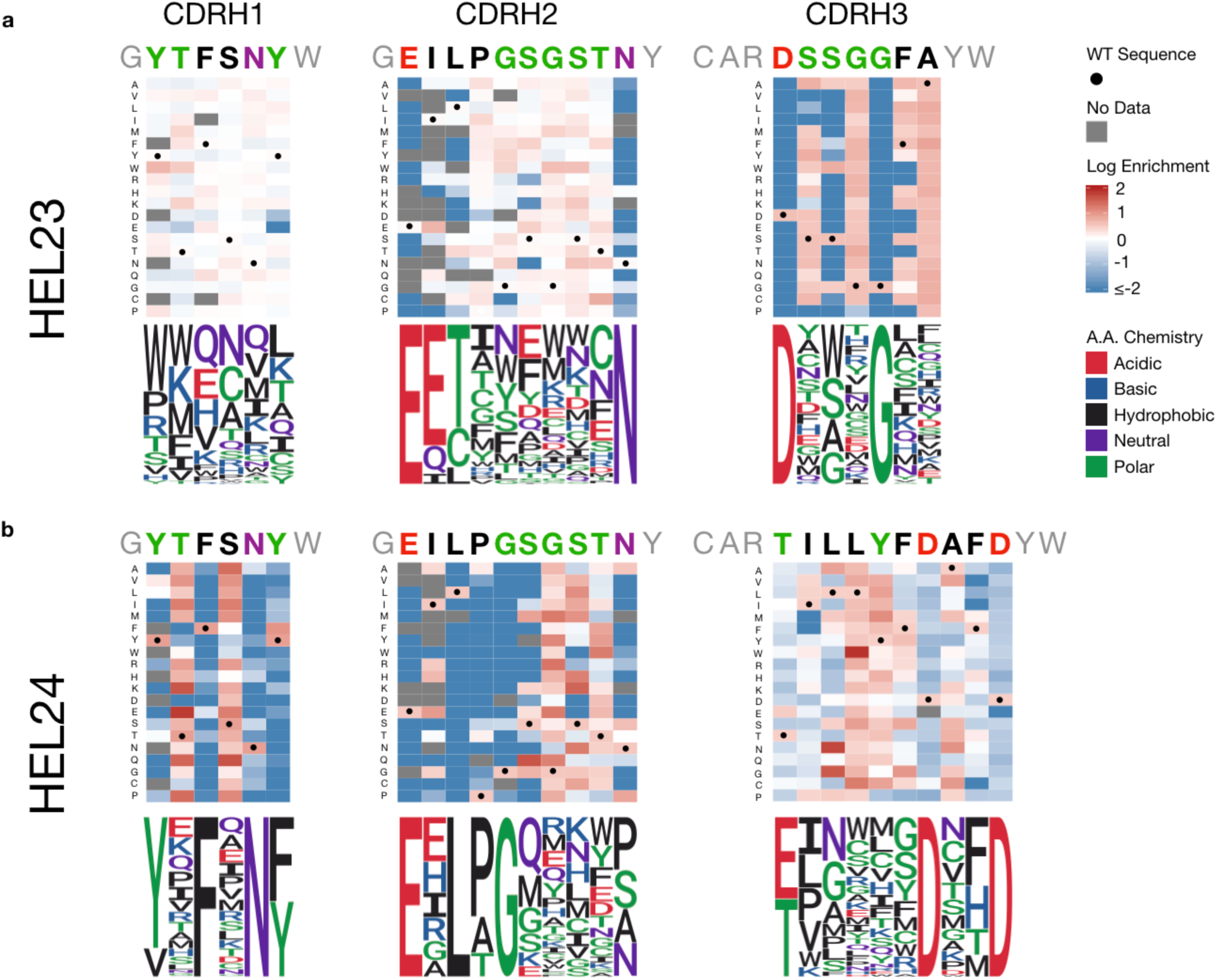
Determining antigen-specificity sequence landscapes by deep mutational scanning (DMS) Heat maps and their corresponding sequence logo plots from the results of DMS of all variable heavy chain CDRs in both the HEL23 (**a**) and HEL24 (**b**) variants reveals positional variance for antigen-binding is clone specific. Libraries of single point-mutations across all heavy chain CDRs were integrated into hybridomas by HDM. NGS was then performed on libraries pre-and post-screening for antigen specificity (**Supplementary Fig. 7, Supplementary Table 3**). Clonal (CDRH3) frequencies of variants post-screening were divided by clonal frequencies of variants pre-screening to calculate enrichment ratios (ERs). Sequence logo plots are generated by normalizing all variants with an ER > 1 (or log[ER] > 0).

Next, by combining the previously acquired knowledge, an HDM library of unique CDRH3 sequences was constructed by transfecting 10^7^ cells with gRNA and a pool of NRO ssODNs with varying degenerate codon lengths, which corresponded to CDRH3 lengths of 10, 12, 14, 15, 16, 18, 20, 22 a.a., thus mimicking the diversity and length distribution found in naïve repertoires. The ssODNs were pooled to mimic the naïve repertoire length distribution at weight-adjusted ratios to account for the expected decrease in integration efficiencies for longer degenerate regions. Following transfection, cells were allowed to divide for >72 hours in order to reach a total cell count >10^8^ cells, at which point antibody expressing cells (Ab+) were labelled with biotinylated anti-IgH and isolated using magnetic assisted cell sorting (MACS). NGS libraries were prepared from genomic DNA isolated from cells pre-and post-selection of antibody expression. Since the number of cells transfected was scaled up, NGS sequencing depth was correspondingly increased to 5.1 × 10^6^ and 1.4 × 10^6^ reads for pre-and post-selection libraries, respectively, with a ∼94% alignment success (**Supplementary Table 1**). Analysis of the NGS data revealed exceptional agreement between the theoretically predicted and observed a.a. frequencies across all CDRH3 lengths (MSE = 2.95±1.14) (**Fig. 3b**). Furthermore, the observed CDRH3 length distribution recapitulated what is observed in naïve repertoires (**Fig. 3d**). There were 9.9×10^4^ and 1.47×10^5^ unique CDRH3 sequences identified in the samples pre-and post-selection of Ab+ cells, respectively. A possible explanation for this inconsistency can be attributed to undersampling due to genomic DNA, where copy numbers of a given sequence are expectedly low.

### Antibody screening and affinity maturation with HDM libraries

The HDM library described in the previous section was next used for antibody discovery using a directed evolution and high-throughput screening approach. Most recombinant antibody libraries, not derived from an immunized animal, require substantially more diversity than our NRO library in PnP cells (1.47 × 10^5^ unique CDRH3s), hence phage and yeast display are often used because of their increased throughput. In order to compensate for this, we opted to screen our library against the model antigen HEL; this served as a reasonable proof-of-concept because the rest of the antibody scaffold (excluding CDRH3) was derived from a HEL-specific sequence (HEL23)^16^. Thus, even with a relatively small library, we aimed to determine if we could discover antigen-specific variants possessing unique CDRH3 sequences when compared to HEL23. First, we enriched PnP cells from the library by two rounds of MACS, using a biotinylated-HEL and streptavidin conjugated magnetic beads. For each round of MACS, the library was expanded to a minimum of 5×10^7^ cells in order to contain multiple copies per variant. Following MACS, a definitive HEL-specific population was visible by flow cytometry (**Fig. 4a**). A subsequent two rounds of FACS enrichment were then performed, followed by a single-cell sort. Antigen specificity for the monoclonal populations was verified by flow cytometry (**Fig. 4a**) and ELISA (**Supplementary Fig. 6**). Genotyping identified a novel clone (HEL24), which had a unique and longer CDRH3 sequence when compared to the original HEL23 (Levenshtein distance = 8) (**Fig. 4d**).

In addition to novel antibody discovery, directed evolution for affinity maturation is also an important step to engineering antibodies. Thus, we also aimed to demonstrate that HDM could be used to improve antibody affinity towards existing antigen-binding clones. To increase the affinity of the previously known HEL23 and newly discovered HEL24 clones, saturation mutagenesis libraries were generated along the CDRH3: HDM was performed with a pool of ssODNs with a single NNK codon tiled across CDRH3 (**Fig 4b**). Higher affinity variants were then enriched by FACS, where antibody avidity was normalized by simultaneous labelling for IgG surface expression (**Fig. 4c**). Following 1-2 rounds of FACS enrichment, monoclonal populations were isolated and characterized. PnP cells have the advantage of simultaneously surface expressing and secreting IgG^16^, therefore ELISAs were performed on these variants to confirm that they had similar or improved antigen affinity (**Supplementary Fig. 6**).

### Antigen specificity-sequence landscapes uncovered by HDM-mediated DMS

DMS is a new method in protein engineering, which combines directed evolution with NGS to assess the functional impact of mutations across the protein sequence landscape^39,40^. In a typical DMS experiment with antibodies, saturation mutagenesis is performed on a single position at a time, followed by screening for functional antibody expression, and then again for antigen binding (**Supplementary Fig. 7**). NGS is performed along the various screening and selection steps, thus providing substantial insight into sequence-specificity relationships. Because of the need for large libraries and high-throughput screening, DMS has most often been performed using phage or yeast display systems. With the ability to generate and screen libraries in our mammalian display system, we therefore aimed to perform DMS on our HEL23 and HEL24 binding variants. We generated HDM single-position saturation libraries of CDRH1, CDRH2, and CDRH3 (pooled ssODNs for each CDR, separate transfection for each CDR). Each CDR library was then first selected on the basis of antibody expression and then screened for variants that retained binding to antigen. We extracted genomic DNA and NGS was performed on the V_H_ genes of antibody expressing cells at both pre-and post-antigen selection. Sequencing depth of all libraries ranged from ∼215,000 −890,000 reads, with an alignment success of >90% (**Supplementary Table 3**). NGS data was analyzed by determining the enrichment ratio (ER) of each mutant, which was calculated by examining the clonal frequencies between the pre-and post-antigen selection libraries^39^ (*Equation 3*). Heatmaps representing ER data were constructed for each CDR (**Fig. 5**). Variants with ERs greater than 1 were then normalized per position and transformed into the corresponding sequence logo plots. These profiles of DMS data clearly show residues critical or detrimental for antigen binding and others which are more amenable to mutations. For example, in both HEL23 and HEL24 CDRH3 sequences, there are two a.a. positions confined to single residue to maintain antigen binding. Also, interestingly, even though HEL23 and HEL24 have identical CDRH1 and CDRH2 sequences, the DMS profiles show substantial variation and dependencies on different positions and residues. Where all positions along the CDRH1 of HEL23 appear more receptive to mutations, the stringency of certain residues found along the CDRH1 of HEL24 implies a greater influence on and contribution to antigen binding.

## DISCUSSION

Paramount to any directed evolution and protein engineering strategy is the ability to generate sufficiently sized libraries of variant clones. Since we rely here on Cas9 to introduce libraries directly in the genome of mammalian cells, we first aimed to optimize a range of parameters associated with HDR by developing an experimental assay that coupled antibody genotype to phenotype using PnP cells (**Fig 1a)**. This allowed us to evaluate a series of parameters through which we determined constitutive Cas9 expression within the host cell played the most important role in improving HDR efficiency (**Fig 1c**). This is likely due to availability and abundance of Cas9 protein already localized within the nucleus at the time ssODN donors become available for DNA repair. We also resolved the optimal homology arm length, which unexpectedly did not correspond to the longest ssODN tested (200 nt), but rather to an intermediate ssODN length of 120 nt. The precise reason why such a decrease in integration occurs for longer ssODN donors is not known, but it is hypothesized that longer ssODNs are less accessible, or may even interfere with DNA repair proteins^26^ and could also be attributed to a decrease in transfection efficiency. We also determined that chemical modification of the 5’ and 3’ ends of ssODNs with PS bonds led to higher HDR, most likely because they have additional stability and nuclease resistance. Since several studies have recently shown that inhibition of the NHEJ pathway regulator 53BP1 can improve HDR, we also knocked out 53BP1 and saw an additional improvement in HDR, reaching a maximum in this study of ∼36% (**Fig. 1d**). In the future, it may be possible to further improve HDR by incorporating additional techniques such as the use of chemically modified gRNAs^41^ and suppression of other NHEJ molecules (e.g., KU70 and DNA ligase IV)^42^, or by exploiting newly engineered variants of Cas9 or other programmable nucleases (e.g., Cpf1)^43,44^.

With parameters optimized for HDR, we next constructed initial HDM libraries targeting CDRH3 through the incorporation of degenerate codons (NNK and NNB) present between the homology arms of the ssODN donors. Following HDM and NGS analysis, we were able to quantitatively elucidate that the a.a. frequencies present in genomic libraries were almost exactly what would have been predicted based on the degenerate codon scheme (**Fig 2a**). This implies that ssODN sequences that had additional homology through similarity with the original CDRH3 were not selectively integrated at higher frequencies, suggesting that HDM is unbiased. NGS analysis did however reveal that repair of double-stranded breaks via the NHEJ/MMEJ pathways occurred in a non-random manner, resulting in several highly abundant variants that were disproportionately present in the library (**Fig. 2c**). This phenomenon of Cas9 repair bias has been reported previously^36^. While the possibility to use the error-prone NHEJ mechanism to introduce additional mutagenesis in our library could be considered a benefit, as it has even been used for discovering novel variants of cell signaling pathways^45^ and dissecting enhancer regions^46^; in the context of antibody engineering, the presence of highly redundant variants would have a detrimental effect on library distribution and screening^47^. Therefore, we reduced the propensity of these events by using a ‘frameshift-stop’ gRNA target sequence, where following Cas9-cleavage, NHEJ/MMEJ events that would normally result in in-frame antibody sequences instead resulted in antibody sequences with premature stop codons (**Fig. 2b**,**c**). HDM libraries constructed using this frameshift-stop gRNA sequence (PnP-HEL23.FS cells) showed a much more uniform distribution, thus greatly reducing overrepresented variants (**Fig. 2d**). We expect that any future studies that aim to engineer antibodies (or other proteins) in mammalian cells will benefit from careful design of gRNA sequences to ensure that NHEJ/MMEJ-biased repair does not compromise library distribution.

Even with the improved HDM efficiencies observed here, our mammalian cell libraries (10^5^) are still substantially smaller relative to what can be routinely achieved in phage (>10^10^) and yeast (>10^7^)^2^. While scale-up can partially compensate for this (**Supplementary Fig. 3**), the slower growth rate and throughput associated with mammalian cell culture necessitates a strategy to maximize the functional quality of libraries. To this end, we designed ssODNs by rationally selecting degenerate codon schemes to mimic the a.a. frequencies of a particular diversity space, in this instance the naïve repertoire of mice. Although there are other approaches to build genetic diversity which very closely recapitulate the a.a. frequencies of a given repertoire (e.g., trinucleotide phosphoramidate-based oligonucleotide synthesis)^48–51^, these methods are expensive and require sophisticated gene assembly and cloning strategies. In contrast, HDM relies on commercially synthesizable degenerate codon schemes and requires no gene assembly or cloning, thus representing substantial savings in time, effort, and cost for library generation. Furthermore, we have shown that when compared to standard NNK and NNB degenerate codon schemes, the rationally selected codon schemes of ssODNs were able to more closely resemble the a.a. frequencies of the mouse naïve repertoire (**Fig. 3a**,**b**). To generate the NRO library, we used a very minimal approach of a single degenerate codon ssODN (per CDRH3 length), however with advances in the large-scale synthesis of oligonucleotide arrays^52^, ssODN pools of custom defined sequences could be included to generate more precise HDM libraries.

Next, we constructed a larger library, based primarily on increasing the number of cells transfected. Following HDM and NGS analysis, we identified that there was a minimum of 1.47×10^5^ variants (unique CDRH3 a.a. sequences). Due to the fact that samples were prepared for NGS from genomic DNA with low copy numbers per variant, we believe the library size generated was actually in the range of 5×10^5^ variants according to live cell counts post-transfection and flow cytometry data. Subjecting the library to antigen selection by MACS and FACS resulted in the discovery of a novel CDRH3 sequence (HEL24), which had a different length and large edit distance from the original sequence (HEL23). While our mammalian cell screening was successful at identifying a single new antigen-specific CDRH3 variant, we understand that such an approach is not in itself competitive with more robust synthetic antibody libraries screened by phage and yeast display, which often recover panels of unique CDRH3 clones^3,32,33,53^. This is most assuredly due to lower library size and diversity in our mammalian system. Thus, we don’t envision our approach will have a major role in the discovery of novel antibody clones, but rather we expect the main application will for engineering and optimizing antibodies, starting from a suitable lead candidate (which can be from *in vivo* selection or phage and yeast display screening). To this end, we showed that by pooling ssODNs with degenerate codons tiled along the CDRH3, that HDM could be used to rapidly engineer our “lead candidate” HEL23 and HEL24 antibodies for higher affinity (**Fig. 4c**,**d, Supplementary Fig. 6**).

One of the most exciting applications of HDM is the ease at which we can perform DMS, a new technique in protein engineering that uncovers sequence-function relationships. DMS relies on the construction of saturation mutagenesis libraries, which requires PCR mutagenesis or synthetic genes and cloning into plasmid expression vectors, thus most DMS studies have been performed in bacteria, phage, and yeast^25,39,54,55^. One previous example performed DMS on an antibody clone using plasmid transfection in mammalian cells^56^, however, the transient presence and polyclonality of plasmids make this a challenging approach. In the case of HDM, we were able to build saturation mutagenesis libraries for DMS in a similar manner in which we did for affinity maturation, by simply pooling ssODNs with single degenerate codons tiled along CDRH1, 2, 3. Because the library sizes required for DMS scale linearly with the number of target positions to investigate, the total library sizes were easily achievable in our hybridoma cells. Furthermore, because no cloning or plasmid transfection was required, DMS libraries could be built rapidly and easily by HDM and genomic integration ensured cellular monoclonality. Analysis of the NGS data produced by DMS revealed residues both critical and detrimental for functional antibody expression and antigen-specificity (**Fig. 5**). This information could be used to rationally select degenerate codons that mimic the functional sequence landscape, which in turn can be used to produce combinatorial HDM libraries that can be screened for antibodies with improved properties (i.e., affinity, specificity, developability). Finally, while in this study we exclusively focus on antibodies, HDM offers the potential to engineer other valuable biological and cellular therapeutics that rely on mammalian expression, such as chimeric antigen-receptors, T cell receptors, cytokine receptors, and intracellular signaling domains^57–60^.

## METHODS

### Hybridoma cell culture conditions

All PnP hybridoma cell lines were cultivated in high-glucose Dulbecco’s Modified Eagle Medium [(DMEM), Thermo Fisher Scientific (Thermo), 11960-044] supplemented with 10% fetal bovine serum [(FBS), Thermo, 16000-044], 100 U/ml Penicillin/Streptomycin (Thermo, 15140-122), 2 mM Glutamine (Sigma-Aldrich, G7513), 10 mM HEPES buffer (Thermo, 15630-056) and 50 μM 2-mercaptoethanol (Sigma-Aldrich, M3148). All hybridoma cells were maintained in incubators at a temperature of 37 °C and 5% CO_2_. Hybridomas were typically maintained in 6 ml of culture in T-25 flasks (TPP, TPP90026), and passaged every 48/72 hours. All hybridoma cell lines were confirmed annually to be negative for *Mycoplasma* contamination. A list of all PnP cell lines are described in **Supplementary Table 4**.

### Guide RNA and ssODN synthesis

For the optimization of parameters or the incorporation of genetic diversity, ssODNs complementary to the non-target strand were ordered directly from Integrated DNA Technologies (IDT) along with any PCR primers or gRNAs used in this study. A recent study has suggested that ssODNs complementary to the non-target strand, and subsequently also complementary to the gRNA, does not compete for Cas9 binding, but instead anneals to the non-target strand further enhancing HDR events^27^. Modified ssODNs had phosphorothioate (PS) in the terminal three nt positions of the 5’ and 3’ ends. A list of all gRNA, primer, and ssODN donor sequences are provided in **Supplementary Table 5** and **Supplementary Table 6**.

### Hybridoma transfection

PnP hybridoma cells were electroporated with the 4D-Nucleofector™System (Lonza) using the SF Cell Line 4D-Nucleofector^®^ X Kit L (Lonza, V4XC-2024, V4XC-2032) with the program CQ-104. Cells were prepared as follows: cells were isolated and centrifuged at 125 × G for 10 minutes, washed with Opti-MEM^®^ I Reduced Serum Medium (Thermo, 31985-062), and centrifuged again with the same parameters. The cells were finally re-suspended in SF buffer (per kit manufacturer guidelines), after which Cas9 plasmid (PX458), Alt-R Cas9 RNP (IDT), or Alt-R gRNA (IDT) and ssODN donor were added. All experiments performed utilize Cas9 from *Streptococcus pyogenes* (SpCas9). Transfections for optimization of parameters were performed by transfecting 2×10^5^ cells with either 1 μg Cas9 plasmid, 100 pmol Alt-R Cas9 RNP, or 115 pmol Alt-R gRNA and 100 pmol ssODN donor in 20 μl, 16-well Nucleocuvette™ strips. All other transfections up to 5×10^6^ cells were performed in 100 μl single Nucleocuvettes™ and reagents were scaled accordingly. Transfections of 10^7^ cells were performed under identical conditions as transfections for 5×10^6^ cells.

### Flow cytometry analysis and sorting

Flow cytometry-based analysis and cell isolation were performed using the BD LSR Fortessa™ and BD FACS Aria™ III (BD Biosciences), respectively. When labeling was required, cells were washed with PBS, incubated with the labeling antibody or antigen for 30 minutes on ice, protected from light, washed again with PBS and analyzed or sorted. The labeling reagents and working concentrations are described in **Supplementary Table 7.** For cell numbers different from 10^6^, the antibody/antigen amount and incubation volume were adjusted proportionally.

### Cell isolation by MACS

MACS isolation of cells was performed using the OctoMACS™ Separator (Miltenyi, 130-042-109) in combination with MS columns (Miltenyi, 130-042-201) for cell counts up to 2×10^8^ cells. Cells were washed with PBS, incubated with the biotinylated antibody or antigen for 30 minutes on ice, washed twice with PBS, resuspended in PBS and Streptavidin Microbeads (Miltenyi, 130-048-102), and incubated in the refrigerator for 15 minutes. Following incubation, cells were washed with additional PBS, and resuspended in 1 ml PBS. The resuspended cells were added to a pre-rinsed magnetic column, washed twice with 500 μl PBS, once with 500 μl growth media. Lastly, the column was removed from the magnetic separator and cells were flushed directly into a collection plate with 1 ml of growth media.

### Measurement of antibody secretion and affinity by ELISA

Sandwich ELISAs were used to measure the secretion of IgG from hybridoma cell lines. Plates were coated with capture polyclonal antibodies specific for V_k_ light chains (goat anti-mouse, Jackson ImmunoResearch, 115-005-174) concentrated at 4 μg/ml in PBS (Thermo, 10010-015). Plates were then blocked with PBS supplemented with 2% m/v milk (AppliChem, A0830) and 0.05% V/V Tween^**®**^-20 (AppliChem, A1389) (PBSMT). Supernatants from cell culture (10^6^ cells/sample, volume normalized to least concentrated samples) were then serially diluted (at 1:3 ratio) in PBS supplemented with 2% m/v milk (PBSM). After blocking, supernatants and positive controls were incubated for 1 hour at RT or O/N at 4°C, followed by 3 washing steps with PBS supplemented with Tween-20 0.05% V/V (PBST). A secondary HRP-conjugated antibody specific for mouse Fc region was used (goat anti-mouse, Sigma-Aldrich, A2554), concentrated at 1.7 μg/ml in PBSM, followed by 3 wash steps with PBST. ELISA detection was performed using a 1-Step™ Ultra TMB-ELISA Substrate Solution (Thermo, 34028) as the HRP substrate, and the reaction was terminated with H_2_SO_4_ (1M). Absorbance at 450 nm was read with Infinite^®^ 200 PRO NanoQuant (Tecan).

For antigen specificity measurements, plates were coated with purified HEL protein (Sigma-Aldrich, 62971-10G-F) concentrated at 4 μg/ml in PBS. Blocking, washing, and supernatant incubation steps were made analogously to the previously described procedure. A secondary HRP-conjugated antibody was used specific for V_k_ light chain (rat anti-mouse, Abcam, AB99617) concentrated at 0.7 μg/ml. ELISA detection by HRP substrate and absorbance reading was performed as previously stated. ELISA data was analyzed with the software GraphPad Prism.

### Sample preparation for NGS

Sample preparation for NGS was performed similar to the antibody library generation protocol of the primer extension method described previously^35^. Genomic DNA was extracted from 1-5×10^6^ cells using the Purelink™ Genomic DNA Mini Kit (Thermo, K182001). All extracted genomic DNA was subjected to a first PCR step. Amplification was performed using a forward primer binding to the beginning of the VH framework region and a reverse primer specific to the intronic region immediately 3’ of the J segment. PCRs were performed with Q5^®^ High-Fidelity DNA polymerase (NEB, M0491L) in parallel reaction volumes of 50 μl with the following cycle conditions: 98°C for 30 seconds; 16 cycles of 98°C for 10 sec, 70°C for 20 sec, 72°C for 30 sec; final extension 72°C for 1 min; 4°C storage. PCR products were concentrated using DNA Clean and Concentrator (Zymo, D4013) followed by 0.8X SPRIselect (Beckman Coulter, B22318) left-sided size selection. Total PCR1 product was amplified in a PCR2 step, which added extension-specific full-length Illumina adapter sequences to the amplicon library. Individual samples were Illumina-indexed by choosing from 20 different index reverse primers. Cycle conditions were as follows: 98°C for 30 sec; 2 cycles of 98°C for 10 sec, 40°C for 20 sec, 72°C for 1 min; 6 cycles of 98°C for 10 sec, 65°C for 20 sec, 72°C for 1 min; 72°C for 5 min; 4°C storage. PCR2 products were concentrated again with DNA Clean and Concentrator and run on a 1% agarose gel. Bands of appropriate size (∼550bp) were gel-purified using the Zymoclean™ Gel DNA Recovery kit (Zymo, D4008). Concentration of purified libraries were determined by a Nanodrop 2000c spectrophotometer and pooled at concentrations aimed at optimal read return. The quality of the final sequencing pool was verified on a fragment analyzer (Advanced Analytical Technologies) using DNF-473 Standard Sensitivity NGS fragment analysis kit. All samples passing quality control were sequenced. Antibody library pools were sequenced on the Illumina MiSeq platform using the reagent kit v3 (2×300 cycles, paired-end) with 10% PhiX control library. Base call quality of all samples was in the range of a mean Phred score of 34.

### Bioinformatics analysis and graphics

The MiXCR v2.0.3 program was used to perform data pre-processing of raw FASTQ files^61^. Sequences were aligned to a custom germline gene reference database containing the known sequence information of the V and J regions for the variable heavy chain of the HEL23-2A synthetic antibody gene. Clonotype formation by CDRH3 and error correction were performed as described by Bolotin et al^61^. Functional clonotypes were discarded if: 1) a duplicate CDRH3 amino acid sequence arising from MiXCR uncorrected PCR errors, or 2) a clone count equal to one. Downstream analysis was performed using R v3.2.2^62^ and Python v2.7.13^63^. Graphics were generated using the R packages ggplot2^64^, RColorBrewer^65^, and ggseqlogo^66^.

### Codon Selection for Library Design

We aimed at designing a library of immunological relevance by investigating all 3,375 degenerate codon schemes with regard to every possible combination described by the IUPAC nucleotide codes. To this end, the a.a. frequencies of each degenerate codon scheme were calculated by dividing the number of codons that encode for a specific a.a. by the total number of codons encoded for a given degenerate scheme. The a.a. frequencies per position of the CDRH3 found in the murine naïve antibody repertoire were based on NGS datasets of V_H_ genes from sorted naïve B-cells, 433,618 unique CDHR3 sequences from 19 C57BL/6 mice, described previously in Greiff et al.^23^. Due to the canonically high frequencies of ‘CAR’ and ‘YW’ residues at the beginning and end of the CDRH3 (Kabat numbering positions 104-106 and 117-118), these positions were kept constant. Utilizing *Equation 1*, an optimized degenerate codon per a.a. position can be determined by calculating the MSE of each degenerate codon relative to the naïve repertoire and then selecting the scheme with the minimum MSE.

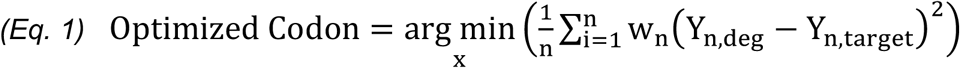

Where x is the degenerate codon, n is the number of a.a. (20), w_n_ is the a.a. weighting factor depending on each a.a.’s frequency in the target, Y_n,deg_ is the a.a. frequency of the degenerate codon, and Y_n,target_ is the a.a. frequency of the target, in this case, the mouse naïve antibody repertoire. To obtain a measure of the overall similarity for the combination of degenerate codons for a CDRH3 of a given length, the mean of each position’s MSE was also calculated.

### Calculating the diversity profiles of libraries

The diversity profile of a given library was calculated as previously described^37^. Briefly, we calculated Hill diversity for *α* = 0 to *α* = 10 by steps of 0.2 according to *Equation 2*,

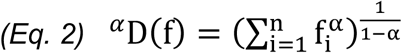

where f_i_ is the clonal frequency of clone i, n is the total number of clones, and *α*-values represent weights. Diversities calculated at alpha values of 0, 1, and 2 represent the species richness, Shannon Diversity, and the exponential inverse Simpson’s Index respectively. Clonal sequences were excluded from the diversity calculation if: 1) a stop codon was present, or 2) the coding sequence was out of frame. A clone was defined based on the exact a.a. sequence of the CDRH3.

### Calculating Levenshtein (edit) distances

From each of the NGS datasets of NRO, NNK, and NNB libraries, 5,000 CDRH3 sequences of length 14 a.a were randomly sampled. Subsequently, for each sampled sequence, the naïve repertoire utilized to develop the NRO degenerate codon scheme was searched for presence of CDRH3 a.a. sequences of Levenshtein (edit) distances 0-6 (**Fig. 3c**).

### Calculation of enrichment ratios (ERs) in DMS

The ERs of a given variant was calculated according to previous methods^39^. Clonal frequencies of variants enriched for antigen specificity by FACS, f_i,Ag+_, were divided by the clonal frequencies of the variants present in the original library, f_i,Ab+_, according to Equation 3.

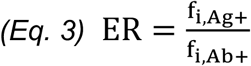

A minimum value of −2 was designated to variants with log[ER] values less than or equal −2 and variants not present in the dataset were disregarded in the calculation. A clone was defined based on the exact a.a. sequence of the CDRH3.

## Data Availability

All cell lines, materials, and relevant data generated in this study are included in this published article (and its supplementary information files) or are available from the corresponding authors upon reasonable request.

We acknowledge the ETH Zurich D-BSSE Single Cell Unit and the ETH Zurich D-BSSE Genomics Facility for support, in particular, T. Lopes, V. Jäggin, E. Burcklen, and C. Beisel. We also thank P. Heuberger for assistance with optimization. This work was supported by the Swiss National Science Foundation (Project #: 31003A_170110 to S.T.R.); the European Research Council Starting Grant (Project #: 679403 to S.T.R.); The National Center of Competence in Research (NCCR) Molecular Systems Engineering (to S.T.R.); the professorship of S.T.R. is supported by an endowment from the S. Leslie Misrock Foundation.

## Author Contributions

D.M.M., C.R.W., and S.T.R. developed the methodology; D.M.M. and S.T.R. designed the experiments and wrote the manuscript; D.M.M., C.P., and S.M.M. performed the experiments; D.M.M. and C.R.W. analyzed sequencing data; C.P., V.G., and W.J.K. provided scientific guidance.

## Supplementary Figure Legends

**Supplementary Fig. 1: Creation of a stable hybridoma cell line with constitutive Cas9 expression**

**a**, A constitutive Cas9 expression cassette contains two genes under control of separate promoters. The first gene encodes for the Cas9-2A-puromycin gene from the plasmid pSpCas9(BB)-2A-Puro (PX459) and enables expression of the Cas9 protein and the puromycin resistance protein from a single transcript. The second gene encodes for the fluorescent protein eGFP used in selection of successfully integrated cassettes. The cassette is integrated into the Rosa26 safe harbor locus of the murine genome by co-transfection with the plasmid pSpCas9(BB)-2A-GFP (PX458). PX458 and PX459 were gifts from Feng Zhang (Addgene plasmid #s 48138/48139). **b**, Validation of Cas9 activity by transfecting the PnP-mRuby hybridoma with only a gRNA complex targeting the mRuby gene. Results from flow cytometry confirms high levels (>90%) of gene knock-out.

**Supplementary Fig. 2: Flow cytometry plots for optimizing HDR parameters**

Flow cytometry plots testing HDR integration efficiencies of all ssODN lengths by transfecting gRNA and modified ssODNs into the Cas9-expressing cell line (PnP-HEL23.FI). 2×10^5^ cells were transfected in replicates and cultured for a minimum of 7 days post-transfection. On day 7, cells were labeled for flow cytometry with a fluorescent antibody (AlexaFluor^®^ 488) targeting the constant region of the antibody heavy chain (IgG2c) and a fluorescent antigen (HEL-AlexaFluor^®^ 647). Data presented is representative of 1 of 2 replicates.

**Supplementary Fig. 3: HDR integration efficiencies when scaling up transfection numbers**

Bar graphs based on flow cytometry measurements of HDR integration efficiencies after scaling up transfection numbers under optimal parameters (Cas9 cell: PnP-HEL23.FI, ssODN length 120 with PS bonds). Cell counts ranging between 10^6^ to 10^7^ were transfected by scaling the amount of reagents accordingly up to 5×10^6^ cells (e.g. 10^6^ cells, 500 pmol gRNA, 500 pmol ssODN donor). The transfection of 10^7^ cells was performed under identical conditions as the transfection for 5×10^6^ cells (e.g. 10^7^ cells, 2.5 nmol gRNA, 2.5 nmol ssODN donor). Cell counts were taken 6 hours post-transfection. 3 days post-transfection, cells were labeled for flow cytometry with a fluorescent antibody (AlexaFluor^®^ 488) targeting the constant region of the antibody heavy chain (IgG2c) and a fluorescent antigen (HEL-AlexaFluor^®^ 647).

**Supplementary Fig. 4: Library design and diversity metrics**

**a**, A comparison between standard randomization schemes, NNK and NNB, and the NRO scheme. For any CDRH3 length (or number of degenerate codons), the NRO scheme has a 0% probability of introducing a nonsense or cysteine mutation. Reducing this probability leads to a higher likelihood of producing a functional antibody sequence following HDM. **b**, Although, there is a reduction in the overall amino acid usage in the NRO scheme, adequate levels of diversity are still maintained, particularly when considering CDRH3 lengths >15, the average length observed in the natural antibody repertoire of mice. Diversity calculations are performed according to the equation above the graph as described by Makowski and Soares^68^ where differences in amino acid frequencies are taken into consideration.

**Supplementary Fig. 5: Impact of degeneracy length on HDM efficiency**

Flow cytometry results for HDM percentages of ssODN donors containing increasing degeneracy/insertion lengths in order to study its impact on integration efficiencies. 2×10^5^ cells were transfected in replicate and cultured for a minimum of 7 days post-transfection. On day 7, cells were labeled for flow cytometry with a fluorescent antibody (AlexaFluor^®^ 488) targeting the constant region of the antibody heavy chain (IgG2c) and a fluorescent antigen (HEL-AlexaFluor^®^ 647). Data presented (mean ± sd) is representative of n = 2.

**Supplementary Fig. 6: Antibody secretion and antigen affinity measurements**

ELISA data for antibody secretion and antigen (HEL) affinity for the original CDRH3 sequences (HEL23, HEL24) and the point mutation variants isolated for higher affinity by flow cytometry (**Fig. 4c**,**d**). Similar secretion profiles indicate a comparable amount of secreted full-length IgG, while difference in antigen affinity profiles indicates similar or increased antigen affinity for point mutation variants compared to the parent sequence. Data presented (mean ± sd) is representative of n = 2 replicates.

**Supplementary Fig. 7: Flow cytometry gating to perform DMS**

Flow cytometry plot displaying an example gating strategy to sort cell populations utilized in DMS experiments. 10^6^ cells are transfected with a pool of ssODNs containing point mutations tiling along the entire CDRH3 sequence. Antibody positive (Ab+) cells are isolated as the control library for NGS and antigen positive (Ag+) cells are isolated as the enriched library for NGS. Samples were prepared for sequencing according to the protocol provided in the **Online Methods** section (**Supplementary Table 3**).

**Supplementary Table 1: Next generation sequencing (NGS) statistics**

NGS statistics for all samples referenced in this study. All sequencing libraries prepared for this study yielded high read counts sufficient for adequate sequencing depth along with a high percentage of alignment (>93%).

**Supplementary Table 2: Overlap analysis between pilot libraries**

Overlap analysis between all 8 NNK/NNB library NGS datasets (**Supplementary Table 1**) indicates that each library generated by HDM produces a unique set of CDRH3 sequences.

**Supplementary Table 3: NGS statistics for DMS studies**

NGS statistics for all DMS samples referenced in this study. All sequencing libraries prepared for this study yielded high read counts sufficient for adequate sequencing depth along with a high percentage of alignment (>90%).

**Supplementary Table 4: Cell lines and descriptions**

A summary table providing a brief description of the hybridoma cell lines generated or used in this study.

**Supplementary Table 5: gRNA target and primer sequences**

A summary table providing the nucleotide sequences of all relevant primers, ssODN donors, and gRNA target sites referenced in this study.

**Supplementary Table 6: ssODN donor sequences**

A summary table providing the nucleotide sequences of all relevant ssODN donors referenced in this study.

**Supplementary Table 7: Flow cytometry labeling concentrations**

A summary table providing information on the fluorescently labeled antibodies and antigens referenced in this study along with their working concentrations.

## References

1. Jain, T. et al. Biophysical properties of the clinical-stage antibody landscape. Proc. Natl. Acad. Sci. 114, 944–949 (2017).

2. Hoogenboom, H. R. Selecting and screening recombinant antibody libraries. Nat. Biotechnol. 23, 1105–1116 (2005).

3. Feldhaus, M. J. et al. Flow-cytometric isolation of human antibodies from a nonimmune Saccharomyces cerevisiae surface display library. Nat. Biotechnol. 21, 163–170 (2003).

4. Köhler, G. & Milstein, C. Continuous cultures of fused cells secreting antibody of predefined specificity. Nature 256, 495–497 (1975).

5. McCafferty, J., Griffiths, A. D., Winter, G. & Chiswell, D. J. Phage antibodies: filamentous phage displaying antibody variable domains. Nature 348, 552–554 (1990).

6. Hanes, J., Jermutus, L., Weber-Bornhauser, S., Bosshard, H. R. & Plückthun, A. Ribosome display efficiently selects and evolves high-affinity antibodies in vitro from immune libraries. Proc. Natl. Acad. Sci. 95, 14130–14135 (1998).

7. Doerner, A., Rhiel, L., Zielonka, S. & Kolmar, H. Therapeutic antibody engineering by high efficiency cell screening. FEBS Lett. 588, 278–287 (2014).

8. Mazor, Y., Blarcom, T. V., Mabry, R., Iverson, B. L. & Georgiou, G. Isolation of engineered, fulllength antibodies from libraries expressed in Escherichia coli. Nat. Biotechnol. 25, 563–565 (2007).

9. Beerli, R. R. et al. Isolation of human monoclonal antibodies by mammalian cell display. Proc. Natl. Acad. Sci. 105, 14336–14341 (2008).

10. Waldmeier, L. et al. Transpo-mAb display: Transposition-mediated B cell display and functional screening of full-length IgG antibody libraries. mAbs 8, 726–740 (2016).

11. Bowers, P. M. et al. Coupling mammalian cell surface display with somatic hypermutation for the discovery and maturation of human antibodies. Proc. Natl. Acad. Sci. 108, 20455–20460 (2011).

12. Black, J. B., Perez-Pinera, P. & Gersbach, C. A. Mammalian Synthetic Biology: Engineering Biological Systems. Annu. Rev. Biomed. Eng. 19, 249–277 (2017).

13. Findlay, G. M., Boyle, E. A., Hause, R. J., Klein, J. C. & Shendure, J. Saturation editing of genomic regions by multiplex homology-directed repair. Nature 513, 120 (2014).

14. Ma, L. et al. CRISPR-Cas9–mediated saturated mutagenesis screen predicts clinical drug resistance with improved accuracy. Proc. Natl. Acad. Sci. 114, 11751–11756 (2017).

15. Matreyek, K. A., Stephany, J. J. & Fowler, D. M. A platform for functional assessment of large variant libraries in mammalian cells. Nucleic Acids Res. 45, e102–e102 (2017).

16. Pogson, M., Parola, C., Kelton, W. J., Heuberger, P. & Reddy, S. T. Immunogenomic engineering of a plug-and-(dis)play hybridoma platform. Nat. Commun. 7, 12535 (2016).

17. Chen, F. et al. High-frequency genome editing using ssDNA oligonucleotides with zinc-finger nucleases. Nat. Methods 8, 753–755 (2011).

18. Ran, F. A. et al. Genome engineering using the CRISPR-Cas9 system. Nat. Protoc. 8, 2281–2308 (2013).

19. Renaud, J.-B. et al. Improved Genome Editing Efficiency and Flexibility Using Modified Oligonucleotides with TALEN and CRISPR-Cas9 Nucleases. Cell Rep. 14, 2263–2272 (2016).

20. Li, H. et al. Design and specificity of long ssDNA donors for CRISPR-based knock-in. bioRxiv 178905 (2017). doi:10.1101/178905

21. Yoshimi, K. et al. ssODN-mediated knock-in with CRISPR-Cas for large genomic regions in zygotes. Nat. Commun. 7, 10431 (2016).

22. Mena, M. A. & Daugherty, P. S. Automated design of degenerate codon libraries. Protein Eng. Des. Sel. 18, 559–561 (2005).

23. Greiff, V. et al. Systems Analysis Reveals High Genetic and Antigen-Driven Predetermination of Antibody Repertoires throughout B Cell Development. Cell Rep. 19, 1467–1478 (2017).

24. Fowler, D. M. & Fields, S. Deep mutational scanning: a new style of protein science. Nat. Methods 11, 801–807 (2014).

25. Whitehead, T. A. et al. Optimization of affinity, specificity and function of designed influenza inhibitors using deep sequencing. Nat. Biotechnol. 30, 543 (2012).

26. Yang, L. et al. Optimization of scarless human stem cell genome editing. Nucleic Acids Res. 41, 9049–9061 (2013).

27. Richardson, C. D., Ray, G. J., DeWitt, M. A., Curie, G. L. & Corn, J. E. Enhancing homology-directed genome editing by catalytically active and inactive CRISPR-Cas9 using asymmetric donor DNA. Nat. Biotechnol. 34, 339–344 (2016).

28. Kim, S., Kim, D., Cho, S. W., Kim, J. & Kim, J.-S. Highly efficient RNA-guided genome editing in human cells via delivery of purified Cas9 ribonucleoproteins. Genome Res. 24, 1012–1019 (2014).

29. Platt, R. J. et al. CRISPR-Cas9 Knockin Mice for Genome Editing and Cancer Modeling. Cell 159, 440–455 (2014).

30. Canny, M. D. et al. Inhibition of 53BP1 favors homology-dependent DNA repair and increases CRISPR–Cas9 genome-editing efficiency. Nat. Biotechnol. (2017). doi:10.1038/nbt.4021

31. Fasth, A. et al. Ectopic expression of RAD52 and dn53BP1 improves homology-directed repair during CRISPR–Cas9 genome editing. Nat. Biomed. Eng. 1, 878 (2017).

32. Fellouse, F. A. et al. High-throughput Generation of Synthetic Antibodies from Highly Functional Minimalist Phage-displayed Libraries. J. Mol. Biol. 373, 924–940 (2007).

33. Hoet, R. M. et al. Generation of high-affinity human antibodies by combining donor-derived and synthetic complementarity-determining-region diversity. Nat. Biotechnol. 23, 344–348 (2005).

34. Xu, J. L. & Davis, M. M. Diversity in the CDR3 Region of VH Is Sufficient for Most Antibody Specificities. Immunity 13, 37–45 (2000).

35. Menzel, U. et al. Comprehensive Evaluation and Optimization of Amplicon Library Preparation Methods for High-Throughput Antibody Sequencing. PLoS ONE 9, e96727 (2014).

36. van Overbeek, M. et al. DNA Repair Profiling Reveals Nonrandom Outcomes at Cas9-Mediated Breaks. Mol. Cell 63, 633–646 (2016).

37. Greiff, V. et al. A bioinformatic framework for immune repertoire diversity profiling enables detection of immunological status. Genome Med. 7, (2015).

38. Zhai, W. et al. Synthetic Antibodies Designed on Natural Sequence Landscapes. J. Mol. Biol. 412, 55–71 (2011).

39. Fowler, D. M. et al. High-resolution mapping of protein sequence-function relationships. Nat. Methods 7, 741 (2010).

40. Araya, C. L. & Fowler, D. M. Deep mutational scanning: assessing protein function on a massive scale. Trends Biotechnol. 29, 435–442 (2011).

41. Hendel, A. et al. Chemically modified guide RNAs enhance CRISPR-Cas genome editing in human primary cells. Nat. Biotechnol. 33, 985–989 (2015).

42. Chu, V. T. et al. Increasing the efficiency of homology-directed repair for CRISPR-Cas9-induced precise gene editing in mammalian cells. Nat. Biotechnol. 33, 543–548 (2015).

43. Hu, J. H. et al. Evolved Cas9 variants with broad PAM compatibility and high DNA specificity. Nature (2018). doi:10.1038/nature26155

44. Zetsche, B. et al. Cpf1 Is a Single RNA-Guided Endonuclease of a Class 2 CRISPR-Cas System. Cell 163, 759–771 (2015).

45. Donovan, K. F. et al. Creation of Novel Protein Variants with CRISPR/Cas9-Mediated Mutagenesis: Turning a Screening By-Product into a Discovery Tool. PLOS ONE 12, e0170445 (2017).

46. Canver, M. C. et al. BCL11A enhancer dissection by Cas9-mediated in situ saturating mutagenesis. Nature 527, 192–197 (2015).

47. Glanville, J. et al. Deep sequencing in library selection projects: what insight does it bring? Curr. Opin. Struct. Biol. 33, 146–160 (2015).

48. Knappik, A. et al. Fully synthetic human combinatorial antibody libraries (HuCAL) based on modular consensus frameworks and CDRs randomized with trinucleotides11Edited by I. A. Wilson. J. Mol. Biol. 296, 57–86 (2000).

49. Ashraf, M. et al. ProxiMAX randomization: a new technology for non-degenerate saturation mutagenesis of contiguous codons. Biochem. Soc. Trans. 41, 1189–1194 (2013).

50. Laura Frigotto et al. Codon-Precise, Synthetic, Antibody Fragment Libraries Built Using Automated Hexamer Codon Additions and Validated through Next Generation Sequencing. Antibodies 4, 88–102 (2015).

51. Virnekäs, B. et al. Trinucleotide phosphoramidites: ideal reagents for the synthesis of mixed oligonucleotides for random mutagenesis. Nucleic Acids Res. 22, 5600–5607 (1994).

52. Kosuri, S. & Church, G. M. Large-scale de novo DNA synthesis: technologies and applications. Nat. Methods 11, 499–507 (2014).

53. Lee, C. V. et al. High-affinity Human Antibodies from Phage-displayed Synthetic Fab Libraries with a Single Framework Scaffold. J. Mol. Biol. 340, 1073–1093 (2004).

54. Koenig, P. et al. Mutational landscape of antibody variable domains reveals a switch modulating the interdomain conformational dynamics and antigen binding. Proc. Natl. Acad. Sci. 114, E486–E495 (2017).

55. Wrenbeck, E. E. et al. Plasmid-based one-pot saturation mutagenesis. Nat. Methods 13, 928–930 (2016).

56. Forsyth, C. M. et al. Deep mutational scanning of an antibody against epidermal growth factor receptor using mammalian cell display and massively parallel pyrosequencing. mAbs 5, 523–532 (2013).

57. Sadelain, M., Rivière, I. & Riddell, S. Therapeutic T cell engineering. Nature 545, 423–431 (2017).

58. Spangler, J. B., Moraga, I., Mendoza, J. L. & Garcia, K. C. Insights into Cytokine–Receptor Interactions from Cytokine Engineering. Annu. Rev. Immunol. 33, 139–167 (2015).

59. Roybal, K. T. & Lim, W. A. Synthetic Immunology: Hacking Immune Cells to Expand Their Therapeutic Capabilities. Annu. Rev. Immunol. 35, 229–253 (2017).

60. Kariolis, M. S., Kapur, S. & Cochran, J. R. Beyond antibodies: using biological principles to guide the development of next-generation protein therapeutics. Curr. Opin. Biotechnol. 24, 1072–1077 (2013).

61. Bolotin, D. A. et al. MiXCR: software for comprehensive adaptive immunity profiling. Nat. Methods 12, 380–381 (2015).

62. R. Development Core Team. R: A Language and Environment for Statistical Computing. (R Foundation for Statistical Computing).

63. Van Rossum, G. & Drake, F. L. The Python Language Reference Manual. (Network Theory Ltd., 2011).

64. Wickham, H. ggplot2: Elegant Graphics for Data Analysis. (Springer International Publishing, 2016).

65. Brewer, C. A., Hatchard, G. W. & Harrower, M. A. ColorBrewer in Print: A Catalog of Color Schemes for Maps. Cartogr. Geogr. Inf. Sci. 30, 5–32 (2003).

66. Wagih, O. ggseqlogo: a versatile R package for drawing sequence logos. Bioinformatics 33, 3645–3647 (2017).

67. Doench, J. G. et al. Rational design of highly active sgRNAs for CRISPR-Cas9–mediated gene inactivation. Nat. Biotechnol. 32, 1262–1267 (2014).

68. Makowski, L. & Soares, A. Estimating the diversity of peptide populations from limited sequence data. Bioinformatics 19, 483–489 (2003).

