## Supplementary Materials for "High-throughput antibody engineering in mammalian cells by CRISPR/Cas9-mediated homology-directed mutagenesis"

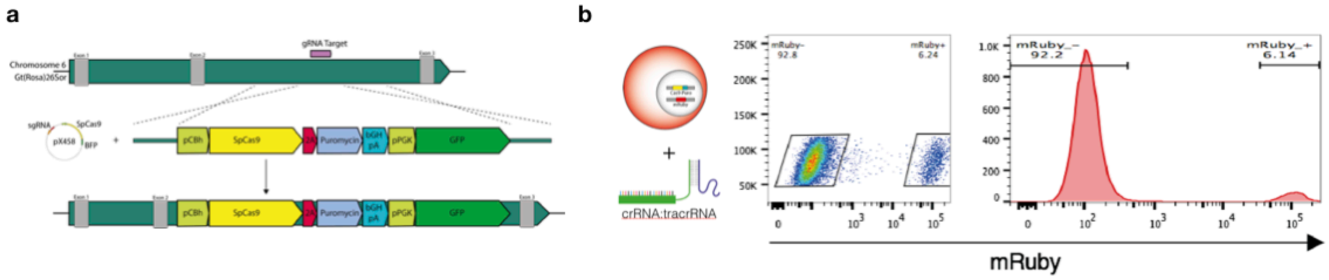


**Supplementary Fig. 1: Creation of a stable hybridoma cell line with constitutive Cas9 expression**

**a,** A constitutive Cas9 expression cassette contains two genes under control of separate promoters. The first gene encodes for the Cas9-2A-puromycin gene from the plasmid pSpCas9(BB)-2A-Puro (PX459) and enables expression of the Cas9 protein and the puromycin resistance protein from a single transcript. The second gene encodes for the fluorescent protein eGFP used in selection of successfully integrated cassettes. The cassette is integrated into the Rosa26 safe harbor locus of the murine genome by co-transfection with the plasmid pSpCas9(BB)-2A-GFP (PX458). PX458 and PX459 were gifts from Feng Zhang (Addgene plasmid #s 48138/48139). **b,** Validation of Cas9 activity by transfecting the PnP-mRuby hybridoma with only a gRNA complex targeting the mRuby gene. Results from flow cytometry confirms high levels (>90%) of gene knock-out.

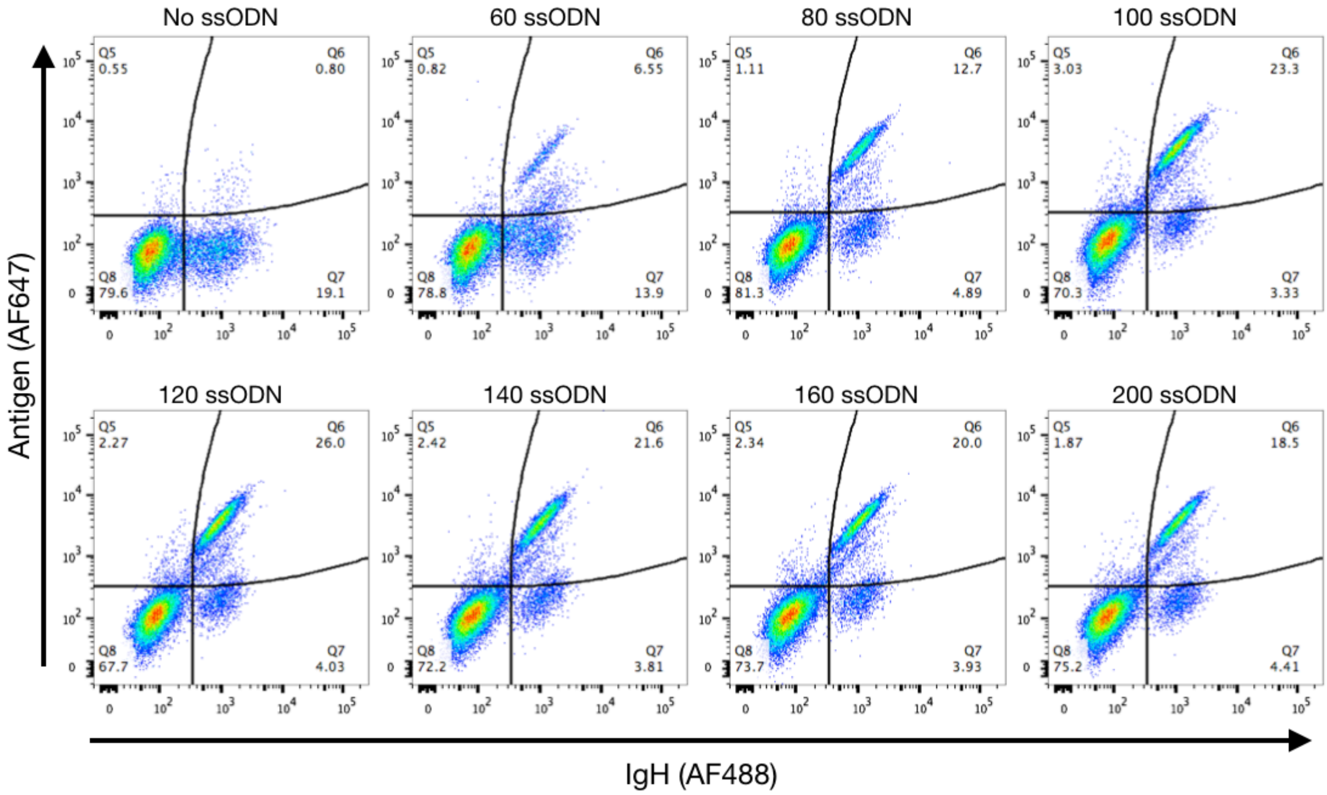


**Supplementary Fig. 2: Flow cytometry plots for optimizing HDR parameters**

Flow cytometry plots testing HDR integration efficiencies of all ssODN lengths by transfecting gRNA and modified ssODNs into the Cas9-expressing cell line (PnP-HEL23.FI). 2x$10^5$ cells were transfected in replicates and cultured for a minimum of 7 days post-transfection. On day 7, cells were labeled for flow cytometry with a fluorescent antibody (AlexaFluor® 488) targeting the constant region of the antibody heavy chain (IgG2c) and a fluorescent antigen (HEL-AlexaFluor® 647). Data presented is representative of 1 of 2 replicates.

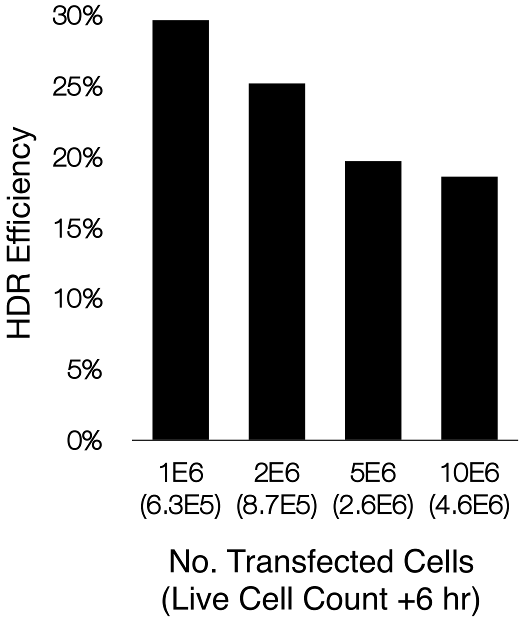


**Supplementary Fig. 3: HDR integration efficiencies when scaling up transfection numbers**

Bar graphs based on flow cytometry measurements of HDR integration efficiencies after scaling up transfection numbers under optimal parameters (Cas9 cell: PnP-HEL23.FI, ssODN length 120 with PS bonds). Cell counts ranging between $10^6$ to $10^7$ were transfected by scaling the amount of reagents accordingly up to $5x10^6$ cells (e.g. $10^6$ cells, 500 pmol gRNA, 500 pmol ssODN donor). The transfection of $10^7$ cells was performed under identical conditions as the transfection for $5x10^6$ cells (e.g. $10^7$ cells, 2.5 nmol gRNA, 2.5 nmol ssODN donor). Cell counts were taken 6 hours post-transfection. 3 days post-transfection, cells were labeled for flow cytometry with a fluorescent antibody (AlexaFluor® 488) targeting the constant region of the antibody heavy chain (IgG2c) and a fluorescent antigen (HEL-AlexaFluor® 647).

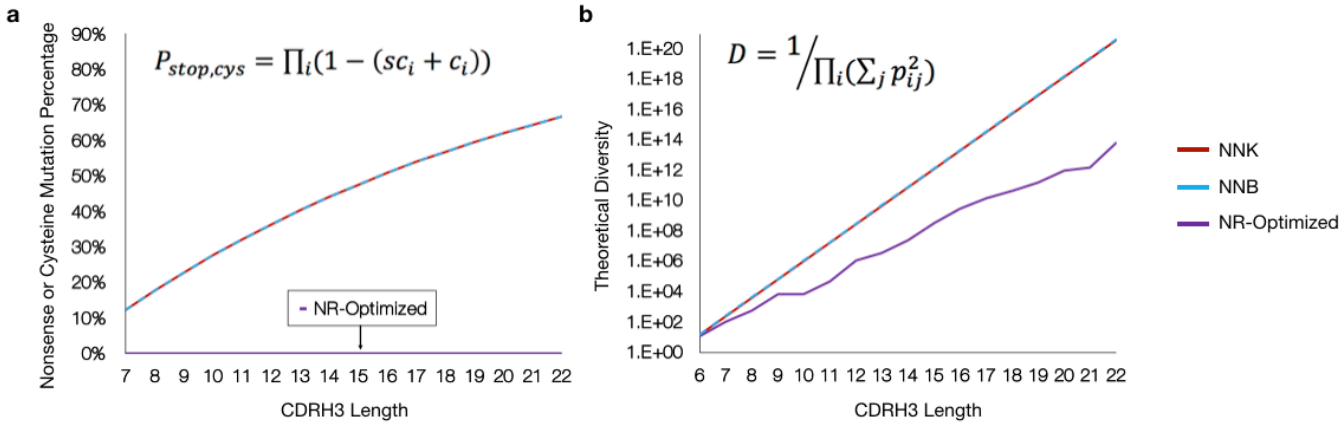


**Supplementary Fig. 4: Library design and diversity metrics**

**a,** A comparison between standard randomization schemes, NNK and NNB, and the NRO scheme. For any CDRH3 length (or number of degenerate codons), the NRO scheme has a 0% probability of introducing a nonsense or cysteine mutation. Reducing this probability leads to a higher likelihood of producing a functional antibody sequence following HDM. **b,** Although, there is a reduction in the overall amino acid usage in the NRO scheme, adequate levels of diversity are still maintained, particularly when considering CDRH3 lengths >15, the average length observed in the natural antibody repertoire of mice. Diversity calculations are performed according to the equation above the graph as described by Makowski and Soares[68] where differences in amino acid frequencies are taken into consideration.

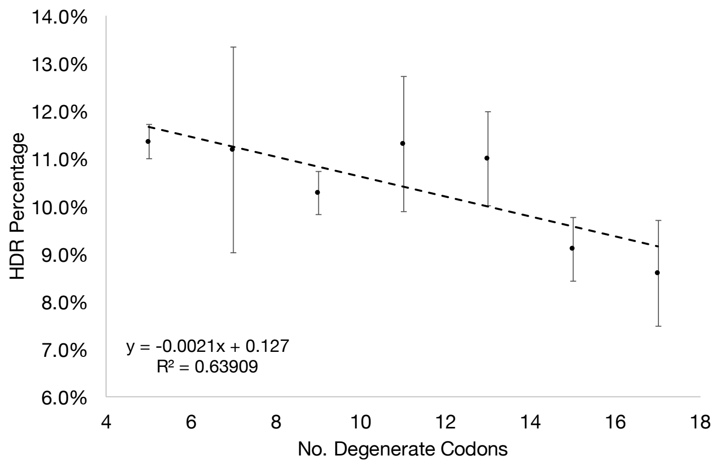


**Supplementary Fig. 5: Impact of degeneracy length on HDM efficiency**

Flow cytometry results for HDM percentages of ssODN donors containing increasing degeneracy/insertion lengths in order to study its impact on integration efficiencies. $2x10^5$ cells were transfected in replicate and cultured for a minimum of 7 days post-transfection. On day 7, cells were labeled for flow cytometry with a fluorescent antibody (AlexaFluor® 488) targeting the constant region of the antibody heavy chain (IgG2c) and a fluorescent antigen (HEL-AlexaFluor® 647). Data presented (mean±sd) is representative of n = 2.

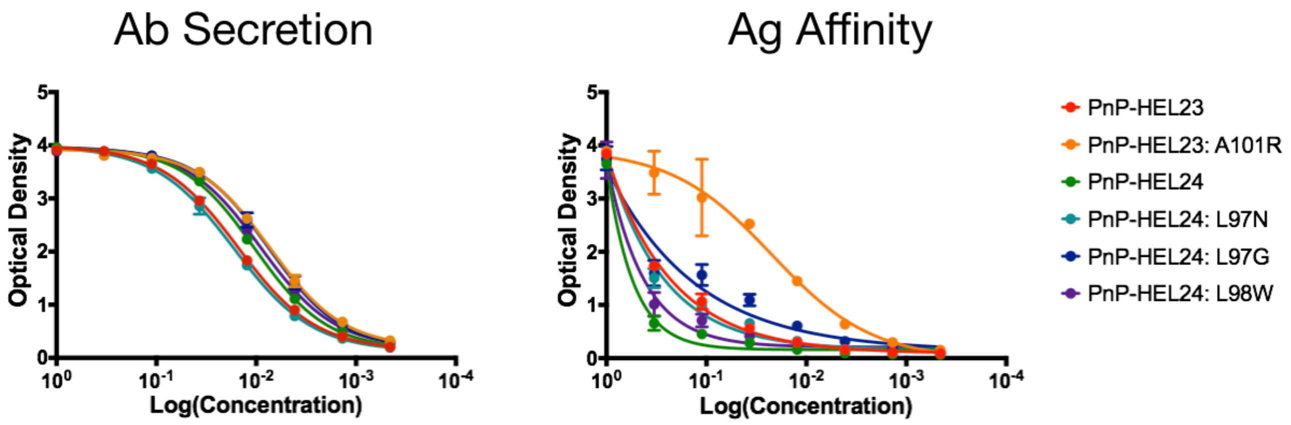


**Supplementary Fig. 6: Antibody secretion and antigen affinity measurements**

ELISA data for antibody secretion and antigen (HEL) affinity for the original CDRH3 sequences (HEL23, HEL24) and the point mutation variants isolated for higher affinity by flow cytometry (**Fig. 4c,d**). Similar secretion profiles indicate a comparable amount of secreted full-length IgG, while difference in antigen affinity profiles indicates similar or increased antigen affinity for point mutation variants compared to the parent sequence. Data presented (mean±sd) is representative of n = 2 replicates.

### HEL23 CDRH3 DMS

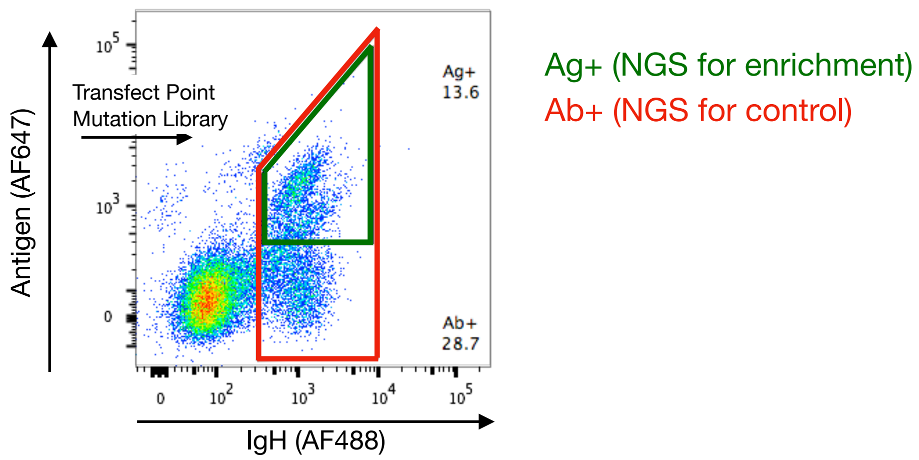


**Supplementary Fig. 7: Flow cytometry gating to perform DMS**

Flow cytometry plot displaying an example gating strategy to sort cell populations utilized in DMS experiments. $10^6$ cells are transfected with a pool of ssODNs containing point mutations tiling along the entire CDRH3 sequence. Antibody positive (Ab+) cells are isolated as the control library for NGS and antigen positive (Ag+) cells are isolated as the enriched library for NGS. Samples were prepared for sequencing according to the protocol provided in the **Online Methods** section (**Supplementary Table 3**).

| Sample | Description | Raw Read count (post-merge) | Aligned Reads | Unique CDRH3s |
|---|---|---|---|---|
| FI-NNK (1) | Pilot library generated by integrating a NNK degenerate codon into the Frameshift-Indel cell line | 577,915 | 551,591 (95.45%) | 13,873 |
| FI-NNK (2) | Pilot library generated by integrating a NNK degenerate codon into the Frameshift-Indel cell line (replicate) | 801,603 | 756,545 (94.38%) | 14,842 |
| FI-NNB (1) | Pilot library generated by integrating a NNB degenerate codon into the Frameshift-Indel cell line | 847,687 | 803,802 (94.82%) | 13,602 |
| FI-NNB (2) | Pilot library generated by integrating a NNB degenerate codon into the Frameshift-Indel cell line (replicate) | 777,070 | 734,095 (94.47%) | 12,773 |
| FS-NNK (1) | Pilot library generated by integrating a NNK degenerate codon into the Frameshift-Stop cell line | 822,301 | 773,378 (94.05%) | 10,329 |
| FS-NNK (2) | Pilot library generated by integrating a NNK degenerate codon into the Frameshift-Stop cell line (replicate) | 810,508 | 759,066 (93.65%) | 10,100 |
| FS-NNB (1) | Pilot library generated by integrating a NNB degenerate codon into the Frameshift-Stop cell line | 780,441 | 729,344 (93.45%) | 8,176 |
| FS-NNB (2) | Pilot library generated by integrating a NNB degenerate codon into the Frameshift-Stop cell line (replicate) | 549,813 | 514,451 (93.57%) | 8,054 |
| NR-O Ab+/- | Library generated by transfecting a pool of different length ssODNs containing NR-Optimized codon schemes. Pre-MACS for antibody expressing cells | 5,082,357 | 4,776,253 (93.98%) | 99,233 |
| NR-O Ab+ | Library generated by transfecting a pool of different length ssODNs containing NR-Optimized codon schemes. Post-MACS for antibody expressing cells | 1,444,914 | 1,364,764 (94.45%) | 146,551 |

**Supplementary Table 1: Next generation sequencing (NGS) statistics**

NGS statistics for all samples referenced in this study. All sequencing libraries prepared for this study yielded high read counts sufficient for adequate sequencing depth along with a high percentage of alignment (>93%).

| Sample | FI-NNK (1) | FI-NNK (2) | FI-NNB (1) | FI-NNB (2) | FS-NNK (1) | FS-NNK (2) | FS-NNB (1) | FS-NNB (2) |
|---|---|---|---|---|---|---|---|---|
| FI-NNK (1) | - | 0 | 0 | 0 | 0 | 0 | 0 | 0 |
| FI-NNK (2) | | - | 0 | 0 | 0 | 0 | 0 | 0 |
| FI-NNB (1) | | | - | 0 | 0 | 0 | 0 | 0 |
| FI-NNB (2) | | | | - | 0 | 1 | 0 | 0 |
| FS-NNK (1) | | | | | - | 0 | 0 | 0 |
| FS-NNK (2) | | | | | | - | 0 | 0 |
| FS-NNB (1) | | | | | | | - | 0 |
| FS-NNB (2) | | | | | | | | - |

**Supplementary Table 2: Overlap analysis between pilot libraries**

Overlap analysis between all 8 NNK/NNB library NGS datasets (**Supplementary Table 1**) indicates that each library generated by HDM produces a unique set of CDRH3 sequences.

| Sample | Description | Raw Read count (post-merge) | Aligned Reads | Point Mutation Variants (Observed/Possible) |
|---|---|---|---|---|
| 23-1-Ab | HEL23 CDRH1 DMS - Control Library | 655,558 | 599,806 (91.5%) | 109/115 |
| 23-1-Ag | HEL23 CDRH1 DMS - Antigen Positive | 277.177 | 251.937 (90.89%) | 109/115 |
| 23-2-Ab | HEL23 HEL24-3 DMS - Control Library | 214,682 | 202,528 (94.34%) | 162/191 |
| 23-2-Ag | HEL23 HEL24-3 DMS - Antigen Positive | 464,073 | 427,205 (92.06%) | 140/191 |
| 23-3-Ab | HEL23 CDRH3 DMS - Control Library | 352,683 | 327,838 (92.96%) | 134/134 |
| 23-3-Ag | HEL23 CDRH3 DMS - Antigen Positive | 784,781 | 721,275 (91.91%) | 130/134 |
| 24-1-Ab | HEL24 CDRH1 DMS - Control Library | 887,680 | 843,578 (98.03%) | 108/115 |
| 24-1-Ag | HEL24 CDRH1 DMS - Antigen Positive | 455,792 | 427,935 (93.89%) | 81/115 |
| 24-2-Ab | HEL24 HEL24-3 DMS - Control Library | 579,301 | 548,357 (94.66%) | 169/191 |
| 24-2-Ag | HEL24 HEL24-3 DMS - Antigen Positive | 500,914 | 450,736 (89.98%) | 96/191 |
| 24-3-Ab | HEL24 CDRH3 DMS - Control Library | 622,195 | 571,173 (91.8%) | 190/191 |
| 24-3-Ag | HEL24 CDRH3 DMS - Antigen Positive | 463,234 | 427,774 (92.35%) | 185/191 |

**Supplementary Table 3: NGS statistics for DMS studies**

NGS statistics for all DMS samples referenced in this study. All sequencing libraries prepared for this study yielded high read counts sufficient for adequate sequencing depth along with a high percentage of alignment (>90%).

| Cell line | Description |
|---|---|
| PnP-HEL23 | PnP-HEL23 cells are immunogenomically engineered to produce antigen specific antibodies targeting hen egg lysozyme (HEL). The light and heavy chains are expressed from a single transcript containing a 2A, self-cleaving peptide. |
| PnP-HEL23.FI | PnP-HEL23.FI cells are a monoclonal population containing a random nucleotide insertion in the CDRH3 after being targeted with CRISPR/Cas9. The random insertion causes a frameshift mutation leading to dysfunctional antibody expression. Final version of this cell line and subsequent clones includes constitutive Cas9 expression. |
| PnP-HEL23.FS | PnP-HEL23.FS cells are a monoclonal population containing a designed frameshift mutation sequence. This designed sequence contains in-frame stop codons on both the 5’ and 3’ sides of the target cleavage site, decreasing the probability of a non-HDR event leading to functional antibody expression. This in turn, results in a more uniformly diverse library. |
| PnP-HEL23.A101R | PnP-HEL23.A101R cells contain a point mutation in the CDRH3 compared to the original HEL23 sequencing. The point mutation increases the antibody’s affinity for its target antigen, hen egg lysozyme. |
| PnP-HEL24 | PnP-HEL24 cells contain the newly identified CDRH3 sequence after subjecting the generated library to the antibody discovery workflow (Figure 4a). These cells produce antigen specific antibodies targeting hen egg lysozyme. |
| PnP-HEL24.L98N | PnP-HEL24.L98N cells contain a point mutation in the CDRH3 compared to the original HEL24 sequencing. The point mutation increases the antibody’s affinity for its target antigen, hen egg lysozyme. |
| PnP-HEL24.L98G | PnP-HEL24.L98G cells contain a point mutation in the CDRH3 compared to the original HEL24 sequencing. The point mutation increases the antibody’s affinity for its target antigen, hen egg lysozyme. |
| PnP-HEL24.L99W | PnP-HEL24.L99W cells contain a point mutation in the CDRH3 compared to the original HEL24 sequencing. The point mutation increases the antibody’s affinity for its target antigen, hen egg lysozyme. |
| PnP-mRuby | PnP-mRuby cells are genetically engineered to produce the fluorescent reporter protein, mRuby, from the heavy chain locus of the hybridoma genome. |

**Supplementary Table 4: Cell lines and descriptions**

A summary table providing a brief description of the hybridoma cell lines generated or used in this study.

| Description | Sequence |
|---|---|
| PnP-HEL23 CDRH3 sgRNA target | TGCGCGCGTGATAGCAGCGGCGG |
| PnP-HEL23.FI CDRH3 sgRNA target | ATTGCGCGCGTGATAGCAGGCGG |
| PnP-HEL23.FS CDRH3 sgRNA target | TTGTGCACGTTAGAGCCAGGCGG |
| Hybridoma ROSA26 intron 2 sgRNA target | AAGCATGTATTGCTTTACGTGGG |
| Hybridoma 53BP1 sgRNA target | GCAGTTGGTGACCACTAACTCGG |
| PnP-mRuby sgRNA target | GTCATGGAAGGTTCGGTCAACGG |
| PnP-HEL CDRH1 sgRNA target | GCTATACCTTTAGCAACTATTGG |
| PnP-HEL CDRH2 sgRNA target | GGCGAAATTCTGCCGGGCAGCGG |
| PnP-HEL VH forward primer for NGS preparation via primer extension (PCR1) | CCCTCCTTTAATTCCCGAAGTGCAGCTGCAGCAG |
| PnP-HEL VH reverse primer for NGS preparation via primer extension (PCR1) | GAGGAGAGAGAGAGAGGACCTGGAGAGGCCATTC |
| TruSeq Universal Adapter forward primer for NGS preparation via primer extension (PCR2) | UNIVERSAL ADAPTER - NNNN CCCTCCTTTAATTCCC |
| TruSeq Index Adapter reverse primer for NGS preparation via primer extension (PCR2) | ADAPTER INDEX X - NNNN GAGGAGAGAGAGAGAG |

**Supplementary Table 5: gRNA target and primer sequences**

A summary table providing the nucleotide sequences of all relevant primers and gRNA target sites referenced in this study.

| Description | Sequence |
|---|---|
| PnP-HEL23.FS ssODN repair template | CATTCTTACCCGCGCTCACGGTCACCAGGGTGCCCTGGCCCCAGTA-<br>CTAAAACTAGCCGCCTGGCTCTA<br>-ACGTGCACAATAATACACCGCGCTATCTTCGCTGGTCAGGCTGCTCAGCTGCA |
| HEL23 CDRH3 ssODN repair template, length 60 nt | TGCCCTGGCCCCAATA-<br>GGCGAAACCACCCGACGAGTCCCTGGCA<br>-CAATAATACACCGCGC |
| HEL23 CDRH3 ssODN repair template, length 80 nt | GTCACCAGGGTGCCCTGGCCCCAATA-<br>GGCGAAACCACCCGACGAGTCCCTGGCA<br>-CAATAATACACCGCGCTATCTTCGCT |
| HEL23 CDRH3 ssODN repair template, length 100 nt | CGCGCTCACGGTCACCAGGGTGCCCTGGCCCCAATA-<br>GGCGAAACCACCCGACGAGTCCCTGGCA<br>-CAATAATACACCGCGCTATCTTCGCTGGTCAGGCTG |
| HEL23 CDRH3 ssODN repair template, length 120 nt | CATTCTTACCCGCGCTCACGGTCACCAGGGTGCCCTGGCCCCAATA-<br>GGCGAAACCACCCGACGAGTCCCTGGCA<br>-CAATAATACACCGCGCTATCTTCGCTGGTCAGGCTGCTCAGCTGCA |
| HEL23 CDRH3 ssODN repair template, length 140 nt | CTGGAGAGGCCATTCTTACCCGCGCTCACGGTCACCAGGGTGCCCTGGCCCCAATA-<br>GGCGAAACCACCCGACGAGTCCCTGGCA<br>-CAATAATACACCGCGCTATCTTCGCTGGTCAGGCTGCTCAGCTGCATATACGCGGT |
| HEL23 CDRH3 ssODN repair template, length 160 nt | AAATAAAGACCTGGAGAGGCCATTCTTACCCGCGCTCACGGTCACCAGGGTGCCCTGGCCCCAATA-<br>GGCGAAACCACCCGACGAGTCCCTGGCA<br>-CAATAATACACCGCGCTATCTTCGCTGGTCAGGCTGCTCAGCTGCATATACGCGGTGTTGCTGCTG |
| HEL23 CDRH3 ssODN repair template, length 200 nt | AACTCCATAACAAAGGTTAAAAATAAAGACCTGGAGAGGCCATTCTTACCCGCGCTCACGGTCACCAG<br>GGTGCCCTGGCCCCAATA-<br>GGCGAAACCACCCGACGAGTCCCTGGCA<br>CAATAATACACCGCGCTATCTTCGCTGGTCAGGCTGCTCAGCTGCATATACGCGGTGTTGCTGCTGGTA<br>TCCGCGGTAAAGGTCGC |
| 5’ homology arm-<br>CDRH1 sequence<br>-3’ homology arm | GGCCATGGCCCGGACGCTGTTTCACCCAGCCAATCCA-<br>ATAGTTTGAAAAGGTATA<br>-GCCGGTCGCTTTGCAGCTAATTTTC |
| Example sequence for single-site mutagenesis. MNN sequence is tiled along the entire CDRH sequence | GGCCATGGCCCGGACGCTGTTTCACCCAGCCAATCCA-<br>**MNN**GTTTGAAAAGGTATA<br>-GCCGGTCGCTTTGCAGCTAATTTTC |
| 5’ homology arm-<br>CDRH2 sequence<br>-3’ homology arm | CTTTGCCTTTAAATTTTTCGTTATA-<br>GTTGGTGCTGCCGGAACCCGGCAGAATTTC<br>-GCCAATCCATTCCAGGCCATGGCCCG |
| Example sequence for single-site mutagenesis. MNN sequence is tiled along the entire CDRH sequence | CTTTGCCTTTAAATTTTTCGTTATA-<br>**MNN**GGTGCTGCCGGAACCCGGCAGAATTTC<br>-GCCAATCCATTCCAGGCCATGGCCCG |
| 5’ homology arm-<br>HEL23 CDRH3 sequence<br>-3’ homology arm | CATTCTTACCCGCGCTCACGGTCACCAGGGTGCCCTGGCCCCAATA-<br>CGCAAAGCCGCCGCTGCTATC<br>-ACGCGCGCAATAATACACCGCGCTATCTTCGCTGGTCAGGCTGCTCAGCTGCA |
| Example sequence for single-site mutagenesis. MNN sequence is tiled along the entire CDRH sequence | CATTCTTACCCGCGCTCACGGTCACCAGGGTGCCCTGGCCCCAATA-<br>**MNN**AAAGCCGCCGCTGCTATC<br>-ACGCGCGCAATAATACACCGCGCTATCTTCGCTGGTCAGGCTGCTCAGCTGCA |
| 5’ homology arm-<br>HEL24 CDRH3 sequence<br>-3’ homology arm | CATTCTTACCCGCGCTCACGGTCACCAGGGTGCCCTGGCCCCAGTA-<br>GTCAAAGGCGTCGAAGTAGAGGAGAATGGT<br>-ACGTGCACAATAATACACCGCGCTATCTTCGCTGGTCAGGCTGCTCAGCTGCA |
| Example sequence for single-site mutagenesis. MNN sequence is tiled along the entire CDRH sequence | CATTCTTACCCGCGCTCACGGTCACCAGGGTGCCCTGGCCCCAGTA-<br>**MNN**AAAGGCGTCGAAGTAGAGGAGAATGGT<br>-ACGTGCACAATAATACACCGCGCTATCTTCGCTGGTCAGGCTGCTCAGCTGCA |

| Description | Sequence |
|---|---|
| 5’ homology arm-<br>Replacement sequence<br>-3’ homology arm | CATTCTTACCCGCGCTCACGGTCACCAGGGTGCCCTGGCCCCAGTA-<br>$(XXX)_X$<br>-ACGTGCACAATAATACACCGCGCTATCTTCGCTGGTCAGGCTGCTCAGCTGCA |
| CDRH3 length 14 a.a. ssODN repair template, NNK scheme | 5’HA-<br>MNNMNNMNNMNNMNNMNNMNNMNNMNN<br>-3’HA |
| CDRH3 length 14 a.a. ssODN repair template, NNB scheme | 5’HA-<br>VNNVNNVNNVNNVNNVNNVNNVNNVNN<br>-3’HA |
| CDRH3 length 10 a.a. ssODN repair template, NRO scheme | 5’HA-<br>GKCVAWVNBVNBVNB<br>-3’HA |
| CDRH3 length 12 a.a. ssODN repair template, NRO scheme | 5’HA-<br>GKCDAAGDVVNBVNBVNBVNB<br>-3’HA |
| CDRH3 length 14 a.a. ssODN repair template, NRO scheme | 5’HA-<br>GKCDAAGKMGDNGDNGDNGDNKYYVNB<br>-3’HA |
| CDRH3 length 15 a.a. ssODN repair template, NRO scheme | 5’HA-<br>GTCDAAGKMGDNGDNGDNGDNGDNVNBVNB<br>-3’HA |
| CDRH3 length 16 a.a. ssODN repair template, NRO scheme | 5’HA-<br>GTCDAAGKMGKNGDVGDNVHYGDHGDNVNBVNB-<br>3’HA |
| CDRH3 length 18 a.a. ssODN repair template, NRO scheme | 5’HA-<br>GTCVAWGKMGKNGDVGDVVHYVHYGKHGKHGDNNNBVNB<br>-3’HA |
| CDRH3 length 20 a.a. ssODN repair template, NRO scheme | 5’HA-<br>GTCSAWGKMGKMGKVGDNGVBVHYVHYGDHGKHGDDGDNBYYDYB<br>-3’HA |
| CDRH3 length 22 a.a. ssODN repair template, NRO scheme | 5’HA-<br>GTCSAWKKCGTAGTMGDNVNBVNYVNYVNYGDHGDHGDDGDNDNBBYYDYB<br>-3’HA |

**Supplementary Table 6: ssODN donor sequences**

A summary table providing the nucleotide sequences of all relevant ssODN donors referenced in this study.

| Target/ Antigen | Working conc. | Dilution from stock | Incubation volume | Fluorophore | Product ID |
|---|---|---|---|---|---|
| IgG2c | 12 µg/ml | 1:100 | 100 µl | AlexaFluor® 488 | 115-545-208 (Jackson ImmunoResearch) |
| IgK | 2.5 µg/ml | 1:80 | 100 µl | Brilliant Violet 421™ | 409511 (BioLegend) |
| Hen egg lysozyme | 0.99 µg/ml | 1:100 | 100 µl | AlexaFluor® 647 | 62971-10G-F (Sigma-Aldrich) |

**Supplementary Table 7: Flow cytometry labeling concentrations**

A summary table providing information on the fluorescently labeled antibodies and antigens referenced in this study along with their working concentrations.
